# Investigating the *Diuraphis noxia*-*Triticum aestivum* interaction using transcriptomics

**DOI:** 10.1101/2024.10.16.618645

**Authors:** Hendrik W. Swiegers, Christine H. Foyer, Anna-Maria Botha

## Abstract

- Aphids often overcome host plant resistance by developing virulence. However, the underlying mechanisms and the precise modifications involved remain poorly understood. Additionally, the molecular response of resistant host plants during compatible (virulent aphid) and incompatible (avirulent aphid) interactions is another area requiring further investigation.
- Here, *Diuraphis noxia* (Russian wheat aphid) biotypes of varying virulence were transferred from susceptible *Triticum aestivum* to a near-isogenic line containing the *Dn7* resistance gene. Transcriptomes of the *D. noxia* biotypes and the resistant host were sequenced before and after the host shift.
- Transcriptomic analysis indicated that the *D. noxia* biotypes had unique responses which were involved in detoxification (e.g. L-xylulose reductase), membrane transport (e.g. Tert-1) and epigenetic regulation. In addition, many transposable elements were rapidly differentially regulated between biotypes and because of the host shift. In the host plant, all biotypes induced jasmonic acid signalling and terpenoid biosynthesis. However, the mono- and diterpenoid pathways were only upregulated following feeding by the biotypes avirulent to *Dn7* and not SAMv2.
- Rapid differential expression of transposons suggests a mechanism of adaptation for clonally reproducing aphids. Furthermore, SAMv2 may possess mechanisms to modulate the defence response of *Dn7*-containing *T. aestivum*.

## Introduction

Aphids (Hemiptera: Aphididae) constitute a globally significant group of insect pests, causing substantial economic losses in agricultural crops. The specialised phloem-feeding behaviour of aphids leads to yield reduction through nutrient depletion or indirect losses from the transmission of plant viruses. *Diuraphis noxia* (Kurdjumov), commonly known as the Russian wheat aphid (**Figure 1**), poses a threat to the production of cereals, primarily *Triticum aestivum* and *Hordeum vulgare* (Botha, 2021; Van Helden *et al*., 2022). The extensive reliance on synthetic chemical insecticides for aphid control has become increasingly challenging due to the evolution of resistance, adverse environmental and human health consequences, and more stringent regulatory measures (Fray *et al*., 2013; Main *et al*., 2018; Chetty-Mhlanga *et al*., 2021; Demortain, 2021).

**Figure 1.**
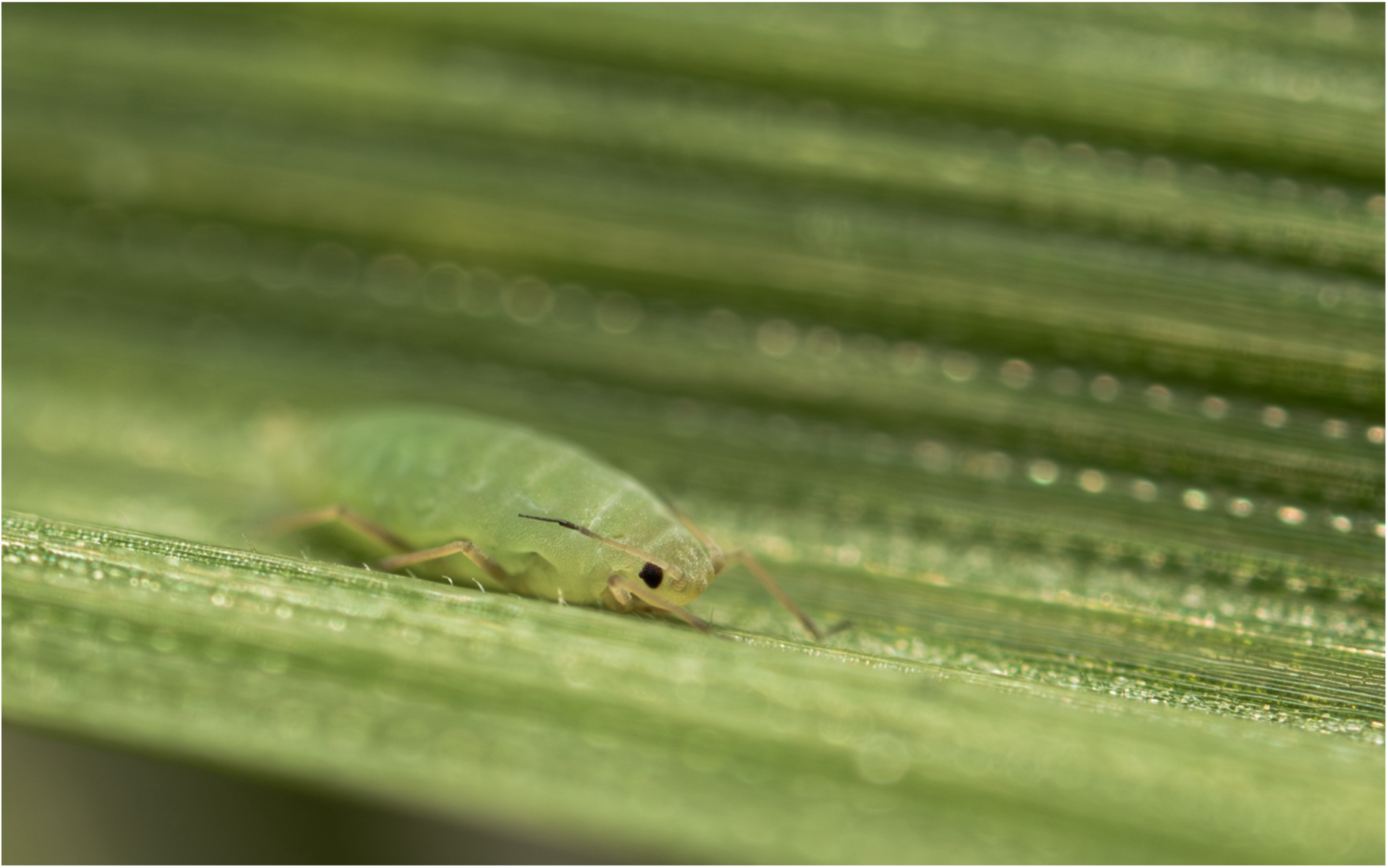
*Diuraphis noxia* (Russian wheat aphid) feeding on *T. aestivum* (bread wheat).

An environmentally safer alternative to manage insect pests comes in the form of host plant resistance (HPR). The exact mechanisms of HPR are often unknown but can be classified as antixenosis (affecting insect host preference), antibiosis (negatively impacting insect survival, growth, and reproduction) or tolerance (the plant’s ability to withstand or recover from insect damage).

Despite the promise of HPR in crop protection, some insects have evolved to overcome HPR after which it is deemed “virulent” against the resistance gene conferring HPR. To date 17 resistance genes (*Dn*) specific to *D. noxia* have been identified in *T. aestivum* (Li *et al*., 2018). Several *D. noxia* populations have in turn developed virulence against many of the *Dn* genes after which it is defined as a new biotype. In South Africa five wild type biotypes (SA1-SA5) have been described with SA1 being the least virulent, while SA5 is the most virulent. Of the *Dn* genes screened, *Dn7* is the only which still confers resistance to all wild type South African *D. noxia* biotypes (Jankielsohn, 2016, 2019). *Dn7* confers resistance to *D. noxia* through antibiosis (Lazzari *et al*., 2009) and antixenosis (Anderson *et al*., 2003; Haley *et al*., 2004) in *T. aestivum*.

Several mechanisms have been described through which aphid pests evolve virulence to HPR. Metabolic resistance involves the upregulation of many gene families to detoxify xenobiotics produced by the host plant (Singh *et al*., 2020). Some level of metabolic resistance will always be required to detoxify the constitutive production of xenobiotics by host plants.

In addition to metabolic resistance, aphids, deliver effectors which modulate the host defence response and nutrient flux (Jones & Dang, 2006). However, when host *R* genes recognise effectors, it results in incompatible interactions. In addition, host recognition of effectors is often biotype specific, indicating that a diverse set of effectors are secreted by different biotypes (Rodriguez & Bos, 2012). This is indicative of the continuous coevolution by aphid and host plant.

As aphids predominantly reproduce through parthenogenesis, genetic variation cannot be obtained through sexual recombination. Therefore, alternative mechanisms must be employed to introduce genetic variability. Such as the transposition of DNA by transposable elements (TEs). TEs are frequently repressed by epigenetic mechanisms to avoid DNA breaks and subsequent deleterious mutations (Horváth *et al*., 2017). However, stress has been shown to remove the repressive epigenetic elements which may include DNA methylation or histone modification (Stapley *et al*., 2015). Thus, activated TEs could facilitate adaptation by generating genetic variation through transposition or TE transcriptional activation through which the expression of nearby genes responsible for adaptation is influenced. The involvement of TEs in aphid adaption to resistant host plants has been observed in *Aphis glycines* and *Myzus persicae* (Singh *et al*., 2020; Yates-Stewart *et al*., 2020). In addition, the virulent *D. noxia* biotype SAM removed DNA methylation by upregulating ten-eleven translocation (TET) methylcytosine dioxygenases (*DnTET)* when fed on resistant *T. aestivum* containing *Dn5* (du Preez *et al*., 2020).

To better understand the formation of virulence in *D. noxia* to resistant *T. aestivum* cultivars, a virulent *D. noxia* biotype, SAMv2 (South African mutant 2) was created in the lab by force-feeding the avirulent biotype SA1 on a *T. aestivum* cultivar containing the *Dn7* resistance gene over a period of 3 months. We investigated the differential transcriptional regulation between SAMv2, its avirulent progenitor, SA1 and the most virulent field biotype, SA5, before and after transferring the aphids from a susceptible *T. aestivum* cultivar to a near-isogenic line containing *Dn7*. In addition, transcriptional regulation of resistant host plant leaves fed on by the aforementioned biotypes were investigated.

## Materials and methods

### Biological material

The *Triticum aestivum* cultivar Gamtoos (*D. noxia* susceptible, denoted “Gamtoos-S”) and its near-isogenic line containing the *D. noxia Dn7* resistance gene (“Gamtoos-R”) were used. Gamtoos-R was developed through a 1RS/1BL translocation from rye to *T. aestivum* (Marais *et al*., 1994; Anderson *et al*., 2003). *T. aestivum* used to maintain aphid colonies or for the host shift experiment (see below) were grown in lightweight expanded clay aggregate at 21ᵒC, 50±10% relative humidity and a photoperiod of 12 h. Plants were watered with modified Hoagland’s solution containing 6.4 mM KNO_3_, 4 mM Ca(NO_3_)2.4H_2_O, 2 mM NH_4_H_2_PO_4_, 2 mM MgSO_4_.7H_2_O, 45 μM FeCl_2_.4H_2_O, 201 μM EDTA, 5 μM SiO_2_, 0.5 μM MnCl_2_.4H_2_O, 0.2 μM ZnSO_4_.7H_2_O, 0.2 μM CuSO_4_.5H_2_O, 4.6 μM H_3_BO_3_ and 0.1 μM (NH_4_)_6_Mo_7_O_24_.4H_2_O.

Colonies of apterous parthenogenetic female aphids of wild type South African *Diuraphis noxia* (Kurdjomov, Aphididae) biotypes SA1 and SA5 (kindly provided by Dr Astrid Jankielsohn, Agricultural Research Council - Small Grain Institute, Bethlehem, South Africa), and the mutant biotype SAMv2 (South African Mutant version 2) were separately established in BugDorm cages (MegaView Science Education Services Co. Ltd.) at 21±2°C and a photoperiod of 12 h. The least and most virulent wild type *D. noxia* biotypes SA1 and SA5 were maintained on Gamtoos-S, while Gamtoos-R was used for SAMv2. SAMv2 was generated in the lab from SA1 by initially feeding it Gamtoos-S, whereafter Gamtoos-R was gradually introduced over a period of three months until the aphids fed exclusively on Gamtoos-R. The virulent SAMv2 and its avirulent progenitor, SA1 can thus be compared directly to study the development of virulence in *D. noxia*.

### Aphid host shift

Gamtoos-S and Gamtoos-R were grown as above for 73 days after which parthenogenic apterous adult *D. noxia* biotypes SA1, SA5 and SAMv2 were placed on Gamtoos-S for 7 days using clip cages. Thereafter, the aphids were transferred to Gamtoos-R for 6 h. Each biological replicate consisted of 10 aphids per plant. For each aphid biotype, four biological replicates were performed (*n* = 4). Samples were taken before (0 h) and after (6 h) the transfer to resistant *T. aestivum* by flash freezing in liquid nitrogen and storing at −80ᵒC. Likewise, three Gamtoos-R biological replicates were sampled before and 6 h after aphid feeding (*n* = 3). *T. aestivum* samples consisted of a 15 mm leaf segment on which aphids were contained. An empty clip cage (ECC) was used as a *T. aestivum* control (*n* = 3).

### RNA extraction and sequencing

Ten aphids (*n* = 10 × 4) were ground to powder with a liquid nitrogen-cooled micropestle after which the RNeasy kit (Qiagen) was used for RNA extraction with on-column DNA digestion (Qiagen), following the manufacturer’s protocol. *T. aestivum* samples were ground to powder using liquid nitrogen pre-cooled mortar and pestles. RNA was extracted from 50 mg of the resulting powder using the RNeasy Plant kit (Qiagen) with on-column DNA digestion (Qiagen), following the manufacturer’s protocol. RNA was precipitated in ethanol before transport to Macrogen for sequencing: 0.1 volume of 3 M sodium acetate (pH 5.5) and 2 volumes of ethanol.

RNA integrity was confirmed with the 2200 TapeStation System (Agilent), while sequencing libraries were constructed with the TruSeq Stranded mRNA Library Prep Kit (Illumina). Paired-end reads of 150 bp in length were sequenced using the NovaSeq 6000 System (Illumina). Four aphid and three plant biological replicates were sequenced, respectively.

### Transcriptome assembly

Quality of raw RNAseq reads were determined using FastQC v0.11.5 (Andrews, 2010) before and after trimming. Trimmomatic v0.39 (Bolger *et al*., 2014) was used to remove sequencing adapters, trimming of the first 7 bases, removal of bases at the start and end with a quality score below 12, trimming of bases with an average quality score below 15 using a sliding window of 10 and to the removal of reads with a length below 36 bp. Using STAR v2.7.10a (Dobin *et al*., 2013), all *D. noxia* reads were aligned to a SAMv2 genome assembled from PacBio HiFi reads (unpublished data; GCA_037042655.1). The alignment was used to generate a genome-guided *D. noxia* transcriptome assembly using Trinity v2.14.0. Similarly, the International Wheat Genome Sequencing Consortium (IWGSC) RefSeq v2.1 genome (Zhu et al., 2021; GenBank assembly: GCA_018294505.1) was used as a guide to generate a *T. aestivum* transcriptome from the Gamtoos-R RNAseq reads. Transcriptome completeness and redundancy were measured with a BUSCO v5.2.2 (Manni *et al*., 2021) analysis using the hemiptera_odb10 or poales_odb10 databases for *D. noxia* and *T. aestivum*, respectively.

### Differential expression and alternative splicing

The *de novo* generated *T. aestivum* transcriptome was compared to that of the NCBI Annotation Release 100 of the IWGSC RefSeq v2.1 genome (Zhu et al., 2021; GenBank assembly: GCA_018294505.1). The latter was used to estimate read counts per transcript using Salmon v1.9.0 (Patro *et al*., 2017) for each *T. aestivum* sample sequenced, while the *de novo* assembled *D. noxia* transcriptome was used in the case of *D. noxia* samples. The following data pre-processing, statistical analysis and generation of volcano plots and heatmaps were performed within the 3D RNA-sep workflow (Guo *et al*., 2021): read counts were scaled to transcript length and library size using the tximport R package (Soneson *et al*., 2015), transcripts were considered expressed if it had a mean count per million above one and was present (read count > 0) in at least 4 or 5 samples for *D. noxia* or *T. aestivum*, respectively. Batch effects were removed using RUVr or RUVs method from the RUVSeq R package (Risso *et al*., 2014) for *D. noxia* or *T. aestivum*, respectively. Data was normalised using the Trimmed Mean of M-values method (Bullard *et al*., 2010). The Limma-voomWithQualityWeights (Law *et al*., 2014; Liu *et al*., 2015; Ritchie *et al*., 2015) function was used to identify differentially expressed (DE) transcripts and genes, differential alternative splicing (DAS; gene level), differential transcript usage (DTU; transcript level).

To assess the effect of feeding on different host plants on the aphid transcriptome, the following contrasts were used for *D. noxia* (Dn): (i) the difference before (0h) and after the host shift (6h) for each biotype: DnSA1_6h - DnSA1_0h, DnSAMv2_6h - DnSAMv2_0h and DnSA5_6h - DnSA5_0h; (ii) the difference before and after the host shift between the average of SA1 and SAMv2: (DnSA1_6h + DnSAMv2_6h)/2 - (DnSA1_0h + DnSAMv2_0h)/2; (iii) difference between biotypes before the host shift: DnSAMv2_0h - DnSA1_0h, DnSA5_0h - DnSA1_0h and DnSA5_0h - DnSAMv2_0h; (iv) difference between biotypes after the host shift: DnSAMv2_6h - DnSA1_6h, DnSA5_6h - DnSA1_6h and DnSA5_6h - DnSAMv2_6h; and lastly (v) the difference in average expression before and after the host shift between SA1 and SAMv2: (DnSAMv2_0h + DnSAMv2_6h)/2 - (DnSA1_0h + DnSA1_6h)/2.

To assess the effect of 6 h-infestation (6h) of the various *D. noxia* biotypes and empty clip cage (ECC) on the host plant (Ta) transcript levels, conditions were compared to the 0 h (0h) control (non-infested host plants, CTRL) using the following contrasts: Ta_ECC_6h - Ta_CTRL_0h, Ta_SA1_6h - Ta_CTRL_0h, Ta_SAMv2_6h - Ta_CTRL_0h and Ta_SA5_6h - Ta_CTRL_0h; gene expression between leaves infested with *D. noxia* biotypes were compared to leaves that had empty clip cages (ECC) clipped to it for 6 h in the contrasts: Ta_SA1_6h - Ta_ECC_6h, Ta_SAMv2_6h - Ta_ECC_6h and Ta_SA5_6h - Ta_ECC_6h; lastly the differences between the containment of the respective *D. noxia* biotypes were determined using the following contrasts: Ta_SAMv2_6h - Ta_SA1_6h, Ta_SAMv2_6h - Ta_SA5_6h and Ta_SA1_6h - Ta_SA5_6h. DAS was calculated by comparing the change of each transcript to the change in gene expression. The various *p-*values are then combined to a single gene-level *p-*value using an F-test. DTU was calculated by comparing the change in expression to the average change in expression of all the remaining transcripts. The false discovery rate method was used to adjust *p-*values for multiple testing (Benjamini & Yekutieli, 2001). An adjusted *p-*value ≥ 0.05 was considered significant, with a log_2_- fold change greater than 1 and a difference in percent spliced (PS; average transcript TPM divided by average gene abundance) greater than 0.1 for differential expression or alternative splicing respectively.

Heatmaps of DE genes were generated by hierarchical clustering using Euclidean distances measurement and Ward’s method (ward.D; Murtagh and Legendre, 2014) using the R functions dist and hclust.

### Gene Ontology and pathway analysis

Gene Ontology (GO) terms and predicted functions of protein products for all differentially expressed *D. noxia* and *T. aestivum* genes were obtained using blastp (Altschup *et al*., 1990) and InterproScan (Quevillon *et al*., 2005) within OmixBox v2.2.4 (BioBam Bioinformatics; Götz et al., 2008). Pathway analyses were performed on differentially expressed *T. aestivum* genes by linking enzyme codes and GO terms to KEGG pathways (Kanehisa *et al*., 2017) and protein blast IDs and GO terms to Plant Reactome pathways (Naithani *et al*., 2020) using OmixBox v2.2.4. Pathway enrichment was performed using Fisher’s exact test and *p-*values adjusted using FDR. All expressed unigenes linked to at least one pathway were used as the reference test.

### Validation of differential gene expression via qPCR

The same RNA samples used for RNAseq, were used for qPCR. cDNA was transcribed from 600 ng RNA (quantified using a NanoDrop 2000, Thermo Scientific) using the SensiFAST cDNA Synthesis Kit, (Meridian Bioscience) following the manufacturer’s protocol. cDNA was quantified fluorometrically using Qubit 4 (Thermo Scientific).

As transcripts from the IWGSC RefSeq v2.1 genome were used for transcript quantification, the corresponding transcript along with its paralogues were firstly identified from the *de novo* assembled Gamtoos-R transcriptome using blastn (Altschup *et al*., 1990). The paralogues were aligned to design primers that where possible, only amplify a single transcript. Primers were designed using primer3 (**Table S1**) (Untergasser *et al*., 2012). For aphid transcripts, primers were designed on the *de novo* assembled *D. noxia* transcriptome (**Table S2**). Reactions for qPCR consisted of 5 μl SensiFAST SYBR no-ROX mix (Meridian Bioscience), 0.5 μl of 10 μM forward and reverse primers (0.4 μl and 0.6 μl for the respective L32 primers; Sapountzis et al., 2014), 0.4 ng template in a total volume of 10 μl. The qPCR reactions were performed using a CFX96 (BioRad) thermocycler set as follows: 95ᵒC for 2 min, 40 cycles of 95ᵒC for 5 s, 10 s at the primer pair’s annealing temperature (**Table S1** and **Table S2**), 72ᵒC for 15 s, followed by a melt curve analysis performed by incremental cooling from 65ᵒC to 95ᵒC. Three technical repeats were performed per sample. Relative expression was calculated using the Pfaffl method (Pfaffl, 2001), taking the reaction efficiency into account for every plate. Reference genes used for *D. noxia* were L27 (Sinha & Smith, 2014) and L32, while 18S and GAPDH were used for *T. aestivum* (Nicolis *et al*., 2017).

Graphs were drawn using Prism v10.0.1 (GraphPad). Significant differences in fold change between experimental conditions were determined with a bootstrapped ANOVA (10,000 iterations) followed by multiple comparisons using the functions t1waybt and mcppb20, respectively, from the R package WRS2 (Mair & Wilcox, 2020).

## Results

To understand the aphid-host plant interaction, virulent (SAMv2) and avirulent (to *Dn7;* SA1 and SA5) *D. noxia* were fed on susceptible Gamtoos-S for one week before the biotypes were transferred to resistant Gamtoos-R and allowed to feed for 6 h. Samples of aphids and plants were taken before and after the host shift from which RNA was extracted and used for RNA sequencing.

### *Diuraphis noxia* transcriptome assembly

Three *D. noxia* biological replicates were sequenced to a depth of ∼20 million reads (∼3 Gb) and a fourth to a depth of ∼60 million reads (∼9 Gb). A *D. noxia* transcriptome was assembled *de novo* using reads from all biotypes investigated (**Table S3**). Of these reads, 72.02% were uniquely mapped while 3.06% were mapped to multiple locations on the biotype SAMv2 genome assembly (GCA_037042655.1), using STAR. The resulting alignment file was provided to Trinity which performed a *de novo* genome-guided transcriptome assembly consisting of 190 000 transcripts and 119 494 unigenes (**Table 1**). A BUSCO analysis on the transcripts using the Hemipteran dataset revealed 99.6% complete (13.5% single copy and 86.1% duplicated), 0.3% fragmented and 0.2% missing universal single-copy orthologs. A unigene-level BUSCO analysis revealed a duplication rate of only 4.6% (**Figure 2**). The *D. noxia* transcriptome was uploaded to the Transcriptome Shotgun Assembly (TSA) database under the accession number: GKVM01000000.

**Figure 2.**
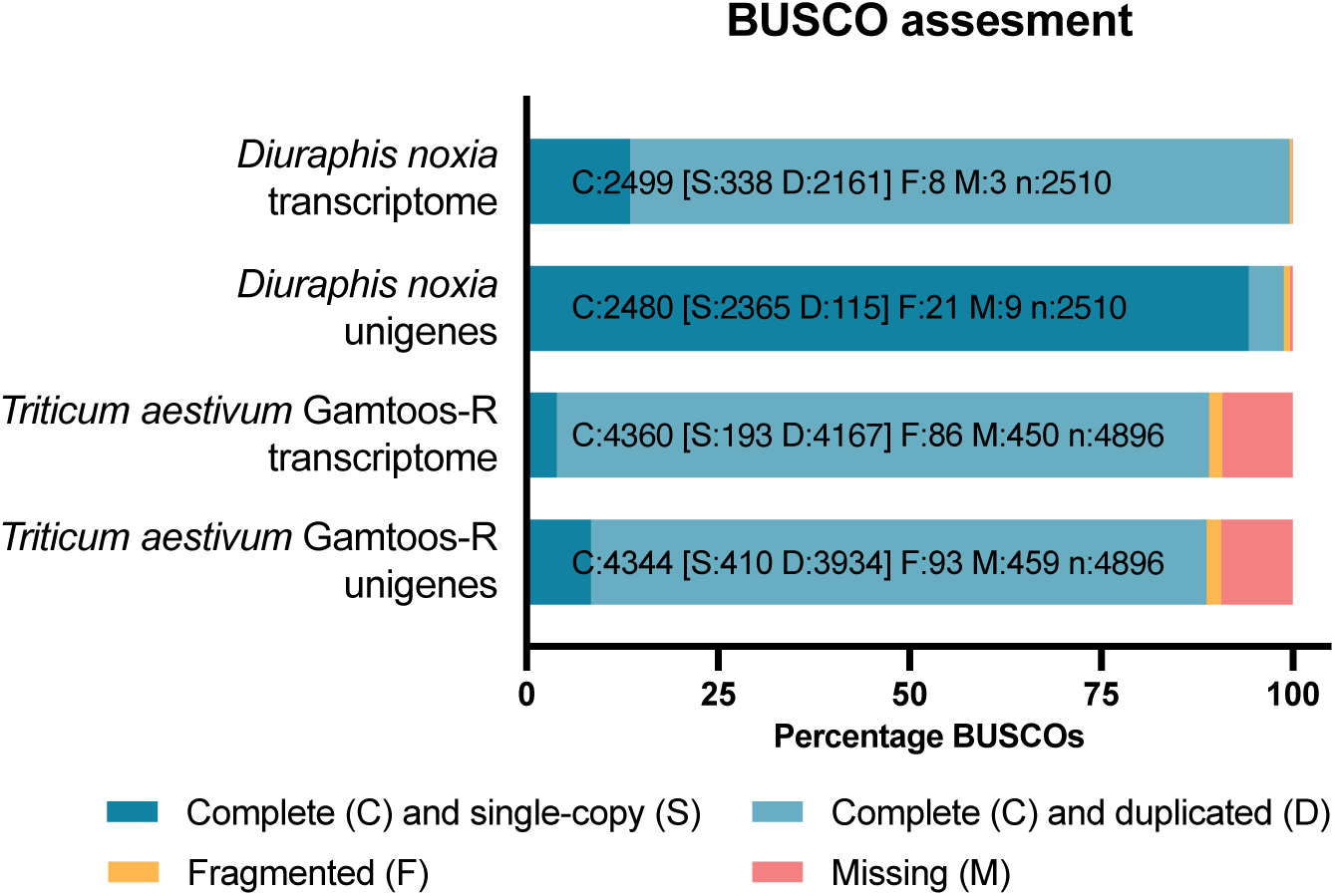
A BUSCO assessment was performed to determine completeness and duplication of the transcriptome assemblies. The universal single-copy orthologs from the hemiptera_odb10 or poales_odb10 databases were used for *Diuraphis noxia* or *Triticum aestivum*, respectively.

**Table 1.** Transcriptome quality parameters for the *Diuraphis noxia* and *Triticum aestivum* Gamtoos-R assemblies. *De novo*, genome-guided assemblies were performed using Trinity v2.14.0. A SAMv2 PacBio HiFi genome assembly or the IWGSC RefSeq v2.1 genome were used as references for *D. noxia* or *T. aestivum*, respectively.

| Quality parameter | *Diuraphis noxia* | *Triticum aestivum* |
| --- | --- | --- |
| # of transcripts | 190 000 | 486,026 |
| Mean length (bp) | 1 313 | 1,270 |
| Median length (bp) | 543 | 798 |
| N50 (bp) | 2 855 | 2,150 |
| %GC | 33.14 | 47.96 |
| # isoforms per unigene | 1.59 | 1.75 |
| # of trimmed reads used | 407 199 392 | 176 309 877 |

### *Triticum aestivum* cultivar Gamtoos-R transcriptome assembly

Likewise, the *T. aestivum* Gamtoos-R reads (**Table S4**) were *de novo* assembled using the IWGSC RefSeq v2.1 genome as guide (**Table 1**, NCBI TSA accession number: GKWO01000000). However, this Gamtoos-R transcriptome had high duplication levels compared to the reference Chinese Spring transcriptome (477 703 vs. 221 416 sequences). This was confirmed by the BUSCO analysis, indicating an 80.4% duplication rate for the unigenes (**Figure 2**). It was therefore decided to proceed with the reference *T. aestivum* transcriptome (IWGSC RefSeq v2.1 NCBI Annotation Release 100) for transcript quantification.

### *Diuraphis noxia* transcriptional regulation

#### *Diuraphis noxia* transcriptional regulation in response to host shift

Transferring *D. noxia* from the susceptible host, Gamtoos-S, to the resistant near isogenic line, Gamtoos-R (containing *Dn7*), resulted in the differential expression of 13 genes in the aphid. Six of which were unique to SA1 and SAMv2 (3 and 3, respectively; **Table 2**). Among those with a predicted protein function, the transmembrane transporters, solute carrier family 26 (SLC26; sulphate transporter) and facilitated trehalose transporter were down regulated (*p =* 0.0083; log_2_FC = −1.14 and *p* = 0.0272; log_2_FC = −1.04, respectively) in SAMv2 after the host shift.

**Table 2.** Number of transcriptional regulation events in *Diuraphis noxia* between various contrasts tested. DE, differential expression; DAS, differential alternative splicing; DTU, differential transcript usage. **Red**, contrasts indicating difference before (0h) and after host shift (6h); **green**, contrasts indicating difference between biotypes before host shift; **orange**, contrasts indicating difference between biotypes after host shift; **blue**, difference average before and after host shift between SA1 and SAMv2. See **Tables S5-S8** for target-specific results.

| Contrast | DE genes | DAS genes | DE transcripts | DTU transcripts |
| --- | --- | --- | --- | --- |
| DnSA1_6h - DnSA1_0h | 3 | 12 | 0 | 12 |
| DnSAMv2_6h - DnSAMv2_0h | 3 | 12 | 0 | 10 |
| DnSA5_6h - DnSA5_0h | 0 | 5 | 0 | 8 |
| (DnSA1_6h + DnSAMv2_6h)/2 - (DnSA1_0h + DnSAMv2_0h)/2 | 9 | 13 | 4 | 22 |
| DnSAMv2_0h - DnSA1_0h | 0 | 9 | 0 | 22 |
| DnSA5_0h - DnSA1_0h | 11 | 16 | 5 | 36 |
| DnSA5_0h - DnSAMv2_0h | 9 | 34 | 3 | 37 |
| DnSAMv2_6h - DnSA1_6h | 19 | 14 | 0 | 22 |
| DnSA5_6h - DnSA1_6h | 7 | 27 | 5 | 41 |
| DnSA5_6h - DnSAMv2_6h | 10 | 39 | 7 | 54 |
| (DnSAMv2_0h + DnSAMv2_6h)/2 - (DnSA1_0h + DnSA1_6h)/2 | 21 | 22 | 0 | 31 |

In addition, two transposons, transposable element P transposase and DDE3 domain containing protein were DE in SAMv2 (*p* = 0.0241; log_2_FC = 3.655 and *p* = 0.0489; log_2_FC = −4.156, respectively). SA1 responded to the host shift by downregulating a predicted venom protease (*p* = 0.0339; log_2_FC = −1.67; **Figure 3**, **Table S5**). All unigenes DE as a result of the host shift are present in clusters 1, 4 and 9 as seen on the heatmap (**Figure 6**).

**Figure 3.**
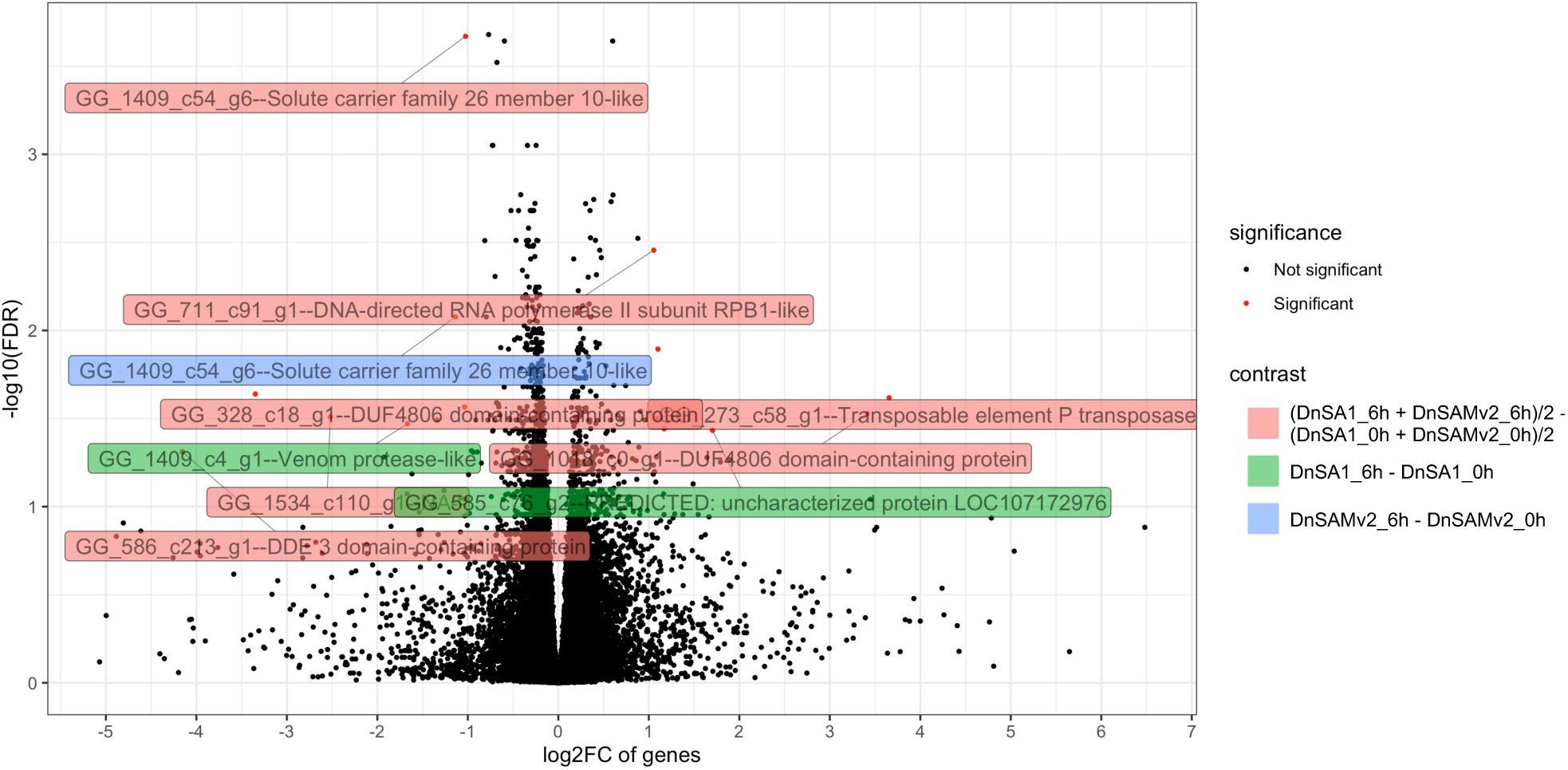
Volcano plot of host shift induced differentially expressed unigenes in *Diuraphis noxia* as a result of the host shift. Contrast groups include: DnSA1_6h - DnSA1_0h, DnSAMv2_6h - DnSAMv2_0h, DnSA5_6h - DnSA5_0h, (DnSA1_6h + DnSAMv2_6h)/2 - (DnSA1_0h + DnSAMv2_0h)/2. Low expression filtered; adjusted *p-*value < 0.05; log2 fold change > 1; labels: gene ID and product description of top 10 distance to 0, 0.

A greater number of differential alternative splicing (DAS) events were present for SA1 (12) and SAMv2 (12; **Table 2**) compared to SA5, with all genes being unique to the respective biotypes.

SA5 was less responsive to the host shift as no significant change in expression was observed in this biotype, however 5 DAS and 8 DTU events were seen. Among these DAS genes was SLC26 (*p =* 0.0344; deltaPS [delta percent spliced] = 0.16) which was also DE and DAS events in SAMv2 after the host shift.

#### *Diuraphis noxia* differential transcriptional regulation between biotypes

No differential expression of unigenes or transcripts were seen between biotype SA1 and its direct descendant, SAMv2 after one week of acclimation to the susceptible host (0h). In contrast, 19 unigenes were differentially expressed between the avirulent and this virulent biotype after transfer to a resistant host. These included unigenes which encode for vasodilator-stimulated phosphoprotein-like, cysteine sulfinic acid decarboxylase, matrix metalloproteinase-16 and a protein containing a reverse transcriptase domain (**Figure 4**, **Table S6**), grouped in clusters 2, 5, 6, 7 and 10 as seen on the heatmap of DE unigenes (**Figure 6**). Alternative splicing was again the preferred mode of transcriptional regulation, and it too increased following the host shift.

**Figure 4.**
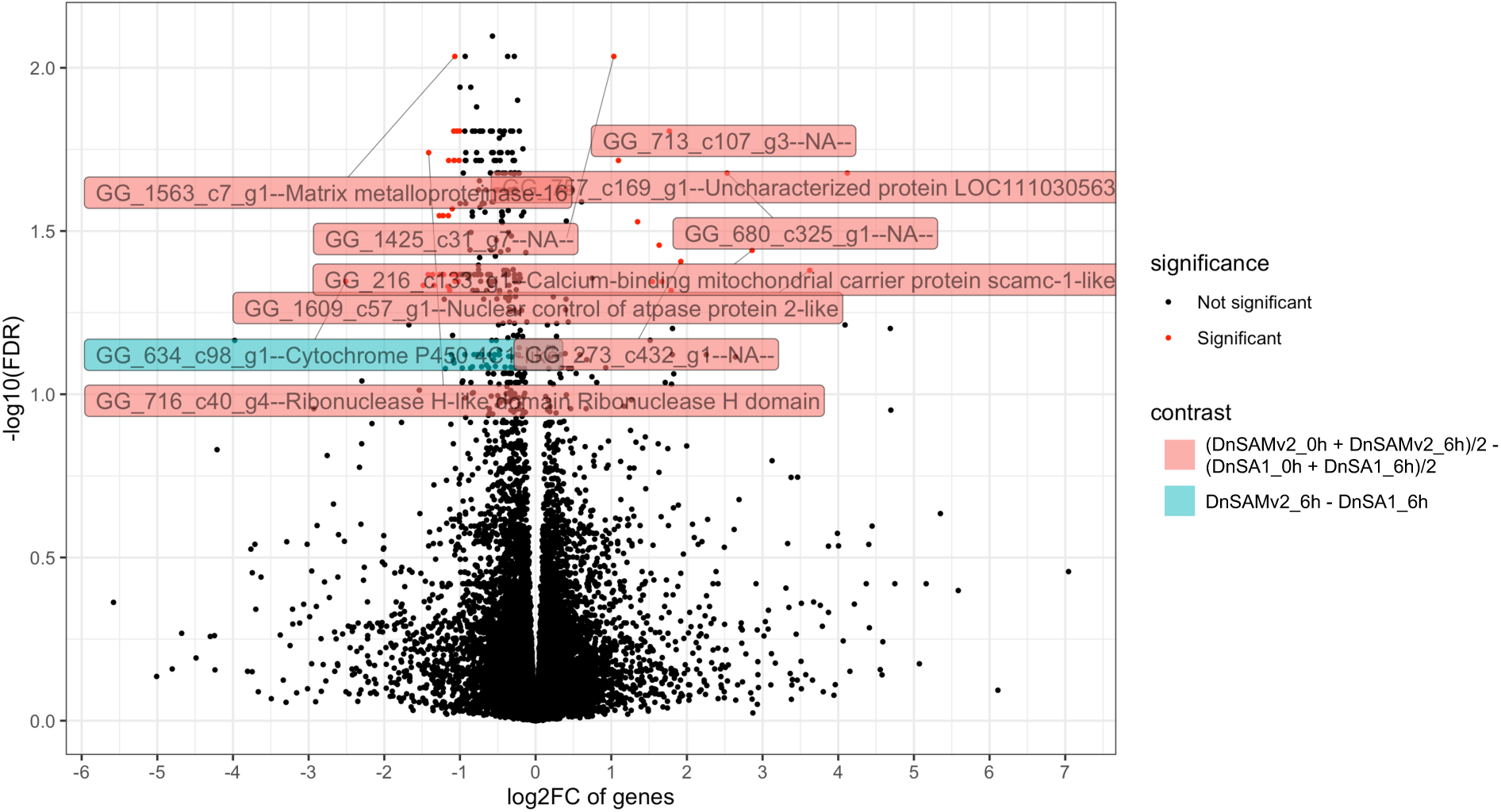
Volcano plot of differentially expressed unigenes between virulent *Diuraphis noxia* biotype SAMv2 and its avirulent progenitor, SA1. Contrasts included: DnSAMv2_0h - DnSA1_0h, DnSAMv2_6h - DnSA1_6h, (DnSAMv2_0h + DnSAMv2_6h)/2 - (DnSA1_0h + DnSA1_6h)/2. Low expression filtered; adjusted *p-*value < 0.05; log2 fold change > 1; labels: gene ID and product description of top 10 distance to 0, 0.

Comparing biotypes SA1 and SAMv2 to SA5 resulted in more transcriptional regulation events than between SA1 and SAMv2. With the only exception being the DE unigenes after the host shift. The most significant differentially expressed unigenes encode for ring canal kelch protein (*kel*), THAP domain-containing protein 1 B-like, Integrase catalytic domain-containing protein, retrovirus-related Pol polyprotein from transposon 17.6 and a protein of unknown function.

*kel* was upregulated between 2.8 and 4.3-fold in SA5 relative to SA1 and SAMv2 (**Table S7**, **Figure 5**). Almost half (7 of 16) of the DE genes between SA5 and the other two biotypes encode for (TEs). These DE unigenes grouped mainly in clusters 3 and 8 on the heatmap (**Figure 6**).

**Figure 5.**
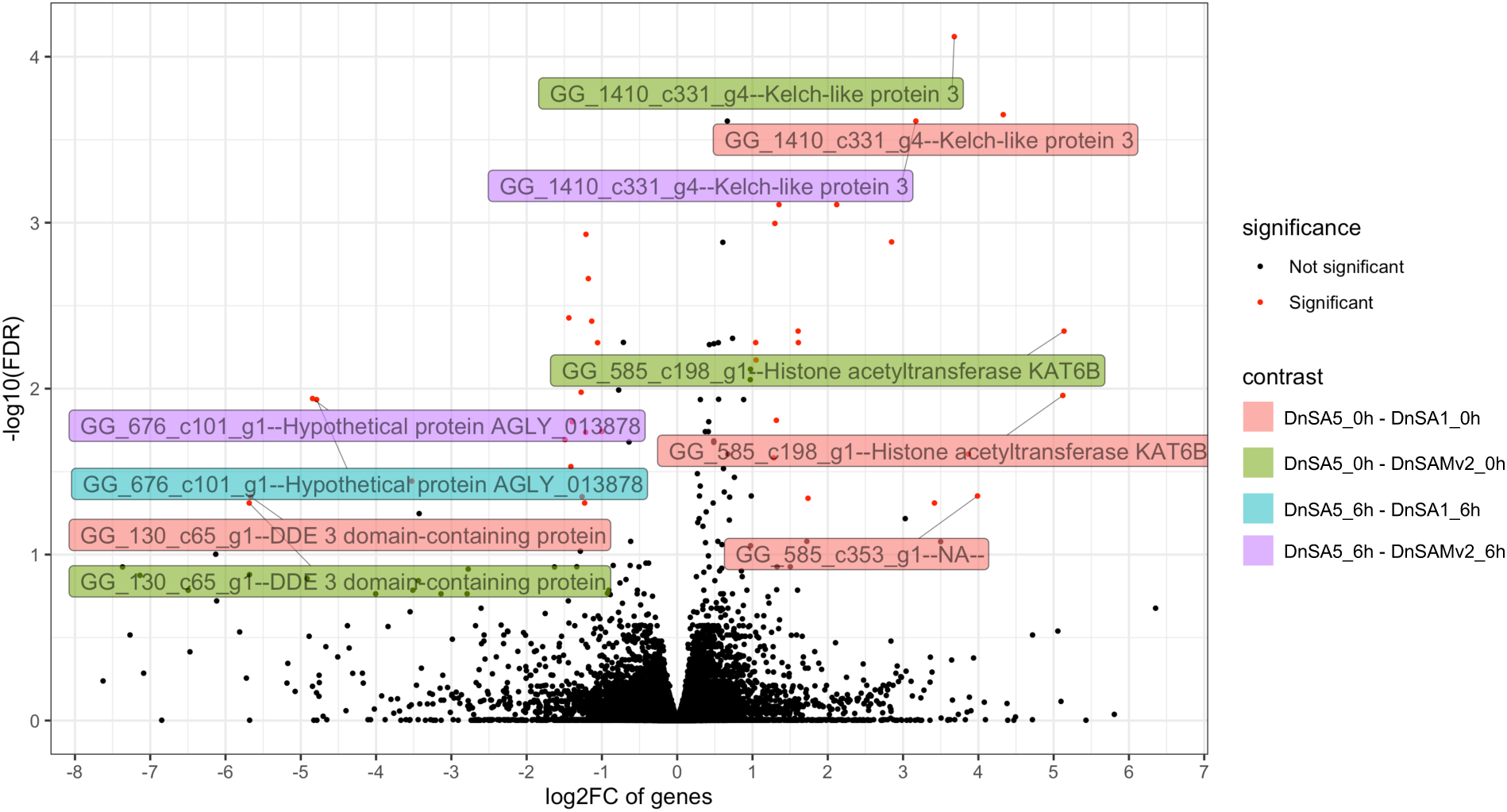
Differentially expressed unigenes between virulent *Diuraphis noxia* biotype SA5 and the biotypes SA1 and SAMv2, respectively. Contrast groups include: DnSA5_0h - DnSA1_0h, DnSA5_0h - DnSAMv2_0h, DnSA5_6h - DnSA1_6h and DnSA5_6h - DnSAMv2_6h. Low expression filtered; adjusted *p-*value < 0.05; log2 fold change > 1; labels: gene ID and product description of top 10 distance to 0, 0.

**Figure 6.**
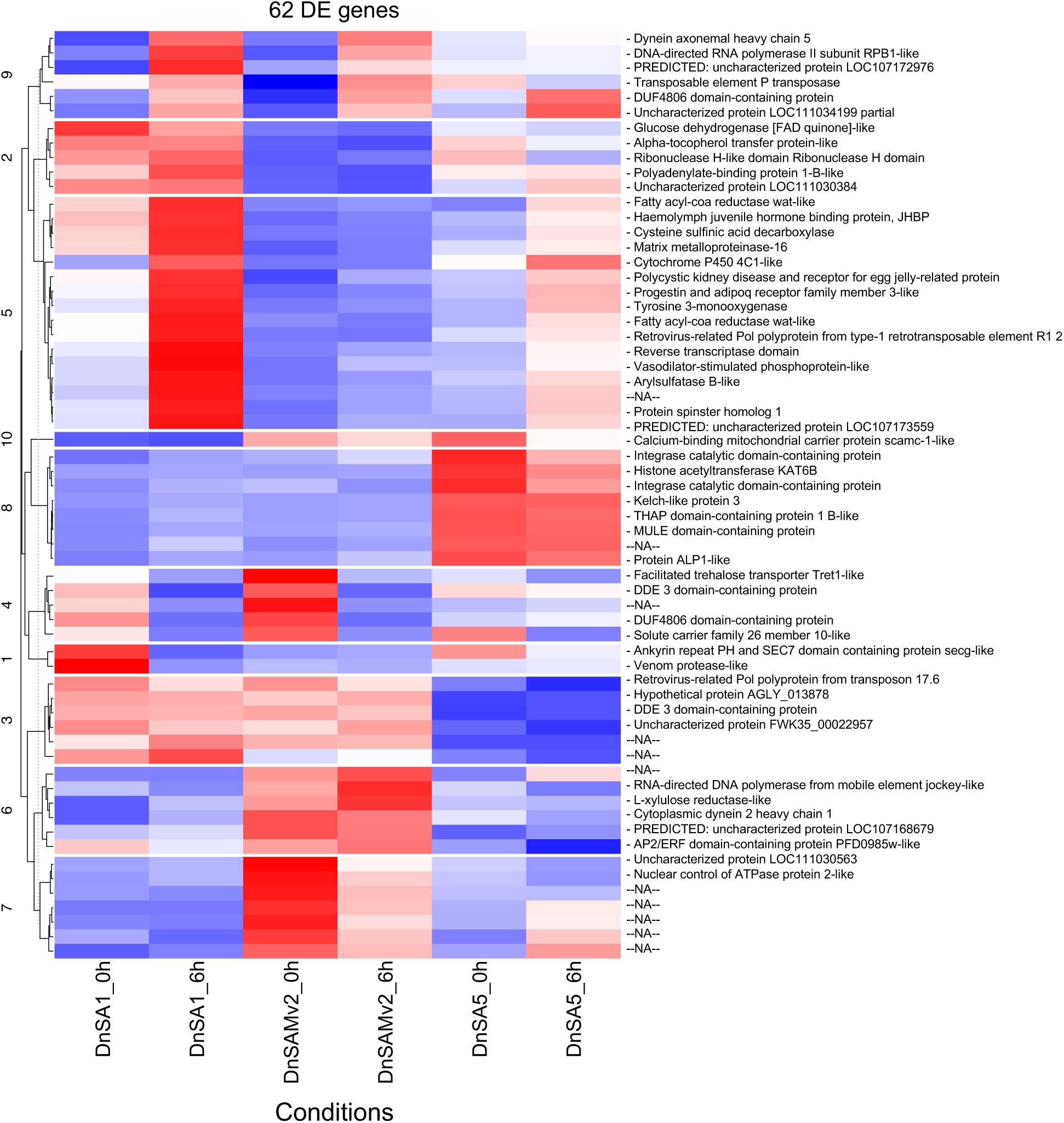
Heatmap demonstrating expression patterns of differentially expressed unigenes in *Diuraphis noxia* biotype SA1, SAMv2 and SA5 before (0h) and 6 h after transferring to resistant *Triticum aestivum* cultivar Gamtoos-R from susceptible cultivar Gamtoos-S. Hierarchical clustering was performed using Euclidean distances measurement and Ward’s method.

### *Triticum aestivum* transcriptional regulation

#### Clip cage-induced *Triticum aestivum* transcriptional regulation

Subsequently, the transcriptional regulation of *T. aestivum* cultivar Gamtoos-R in response to aphid feeding was investigated. Compared to the 0 h control, an empty clip cage placed on the plant for 6 h resulted in the differential expression of 1228 genes (**Figure 7**), which is more than what was found in the host plant tissue that was exposed to feeding of *D. noxia* biotype SA1 in a clip-cage clipped to the plant, but less than the virulent biotypes SAMv2 and SA5 in clip cages clipped to the leaves (**Table 3**). Interestingly, 54% of the clip cage-induced differentially expressed genes were shared with aphid-feeding induced differentially expressed genes (**Figure 8** and **Table S9**). The heatmap of DE *T. aestivum* genes illustrates that the 0 h control is the most distinct (**Figure 10**).

**Figure 7.**
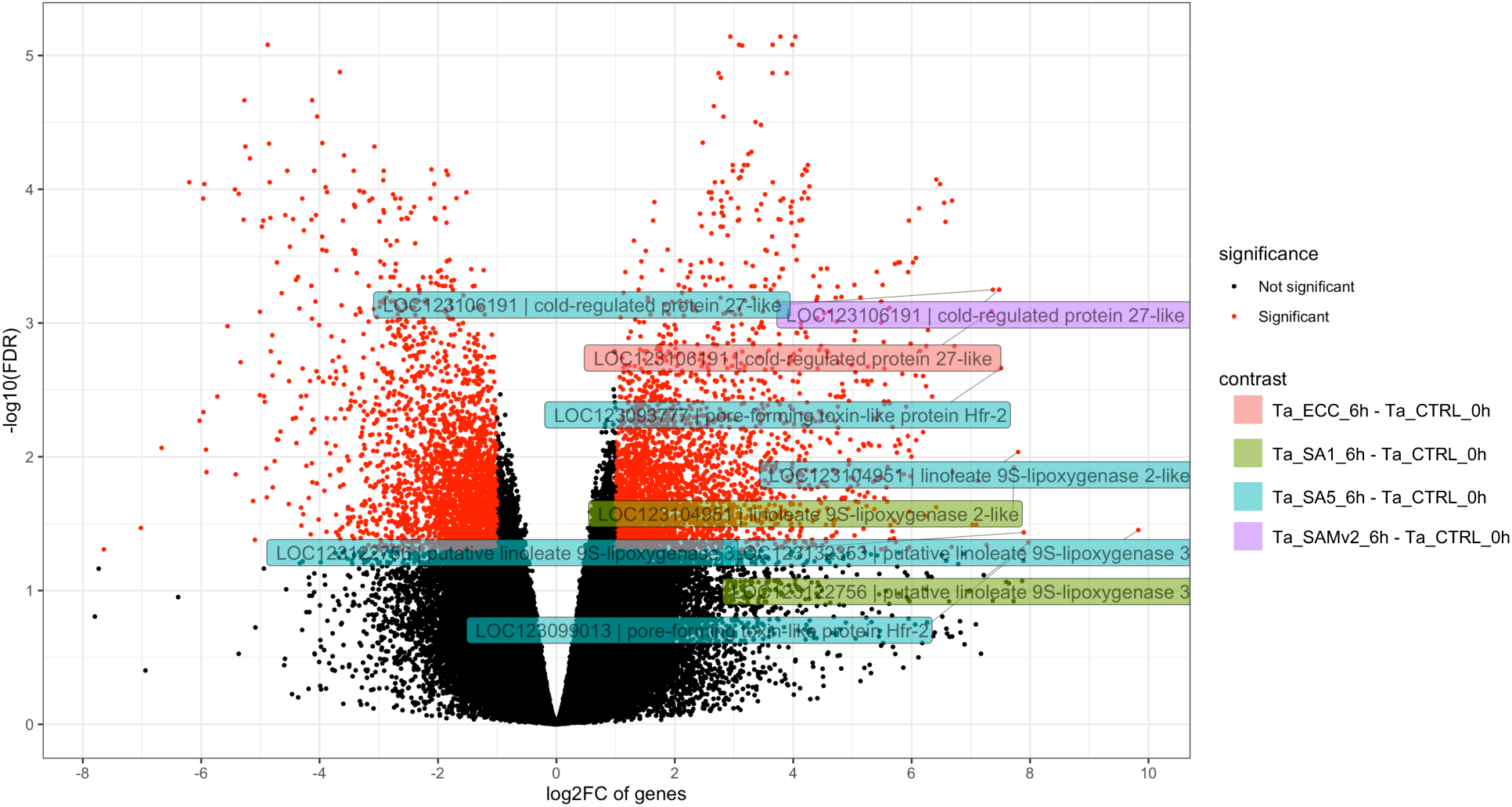
Genes differentially expressed in *Triticum aestivum* Gamtoos-R following the containment of *Diuraphis noxia* biotypes SA1, SAMv2 and SA5 on the plant leaf for 6 h. An empty clip cage (ECC) was used as control. Contrast groups include: Ta_ECC_6h - Ta_CTRL_0h, Ta_SA1_6h - Ta_CTRL_0h, Ta_SAMv2_6h - Ta_CTRL_0h and Ta_SA5_6h - Ta_CTRL_0h. Low expression filtered; adjusted *p*-value < 0.05; log2 fold change > 1; labels: gene ID and product description of top 10 distance to 0, 0.

**Figure 8.**
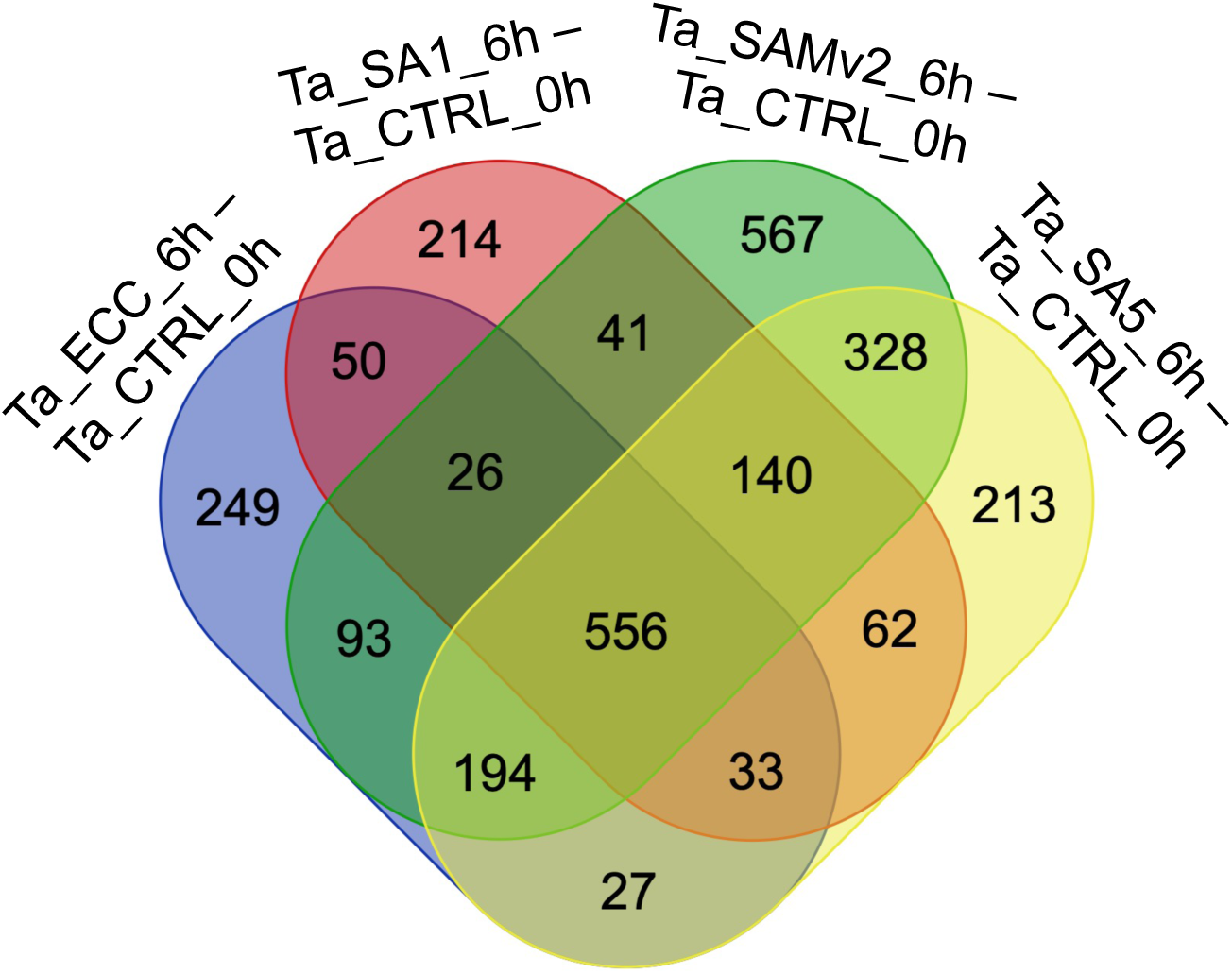
Venn diagram of differentially expressed *Triticum aestivum* genes following the containment of *Diuraphis noxia* on the plant for 6 h using clip cages. An empty clip cage (ECC) was used as control. Contrasts indicating differences between 0 h control and other conditions are included.

**Table 3.** Number of transcriptional regulation events in *Triticum aestivum* following the containment of *Diuraphis noxia* on the plant for 6 h using clip cages. An empty clip cage (ECC) was used as control. DE, differential expression; DAS, differential alternative splicing; DTU, differential transcript usage. **Red**, contrasts indicating differences to the 0h control and other conditions; **green**, contrasts indicating difference between empty clip cage control and *D. noxia* containment; **orange**, contrasts indicating difference between biotypes. See **Tables S9-S11** for target-specific results.

| Contrast | DE genes | DAS genes | DE transcripts | DTU transcripts |
| --- | --- | --- | --- | --- |
| Ta_ECC_6h - Ta_CTRL_0h | 1228 | 59 | 800 | 70 |
| Ta_SA1_6h - Ta_CTRL_0h | 1122 | 77 | 679 | 91 |
| Ta_SAMv2_6h - Ta_CTRL_0h | 1945 | 78 | 1468 | 111 |
| Ta_SA5_6h - Ta_CTRL_0h | 1553 | 72 | 1008 | 99 |
| Ta_SA1_6h - Ta_ECC_6h | 10 | 89 | 1 | 119 |
| Ta_SAMv2_6h - Ta_ECC_6h | 1 | 70 | 0 | 114 |
| Ta_SA5_6h - Ta_ECC_6h | 32 | 74 | 7 | 120 |
| Ta_SAMv2_6h - Ta_SA1_6h | 0 | 116 | 0 | 187 |
| SAMv2v 6h - Ta_SA5_6h | 0 | 108 | 0 | 155 |
| Ta_SA1_6h - Ta_SA5_6h | 0 | 107 | 0 | 152 |

The clip cage also induced DAS in *T. aestivum*, but to a lesser extent compared to DE. These DAS genes encode for a serine/threonine-protein kinase, a pentatricopeptide repeat-containing protein and many uncharacterised proteins (**Table S11**).

#### *Diuraphis noxia*-induced *Triticum aestivum* transcriptional regulation

Comparing the transcripts produced by the host leaves on which empty clip cages were clipped, to leaves on which aphids were contained with clip cages for 6 h, revealed that 32 *T. aestivum* genes were significantly upregulated (**Figure 9**; **Table 3** and **Table S10**). Of these, 17 clustered together (cluster 11 as seen on the heatmap; **Figure 10**) and included the TIFY transcription factors 10C, 10B and 11B, transcription factors bHLH 6 and 148, linoleate 9S-lipoxygenase 3 (LOX1.3) and allene oxide synthase 2 (AOS2).

**Figure 9.**
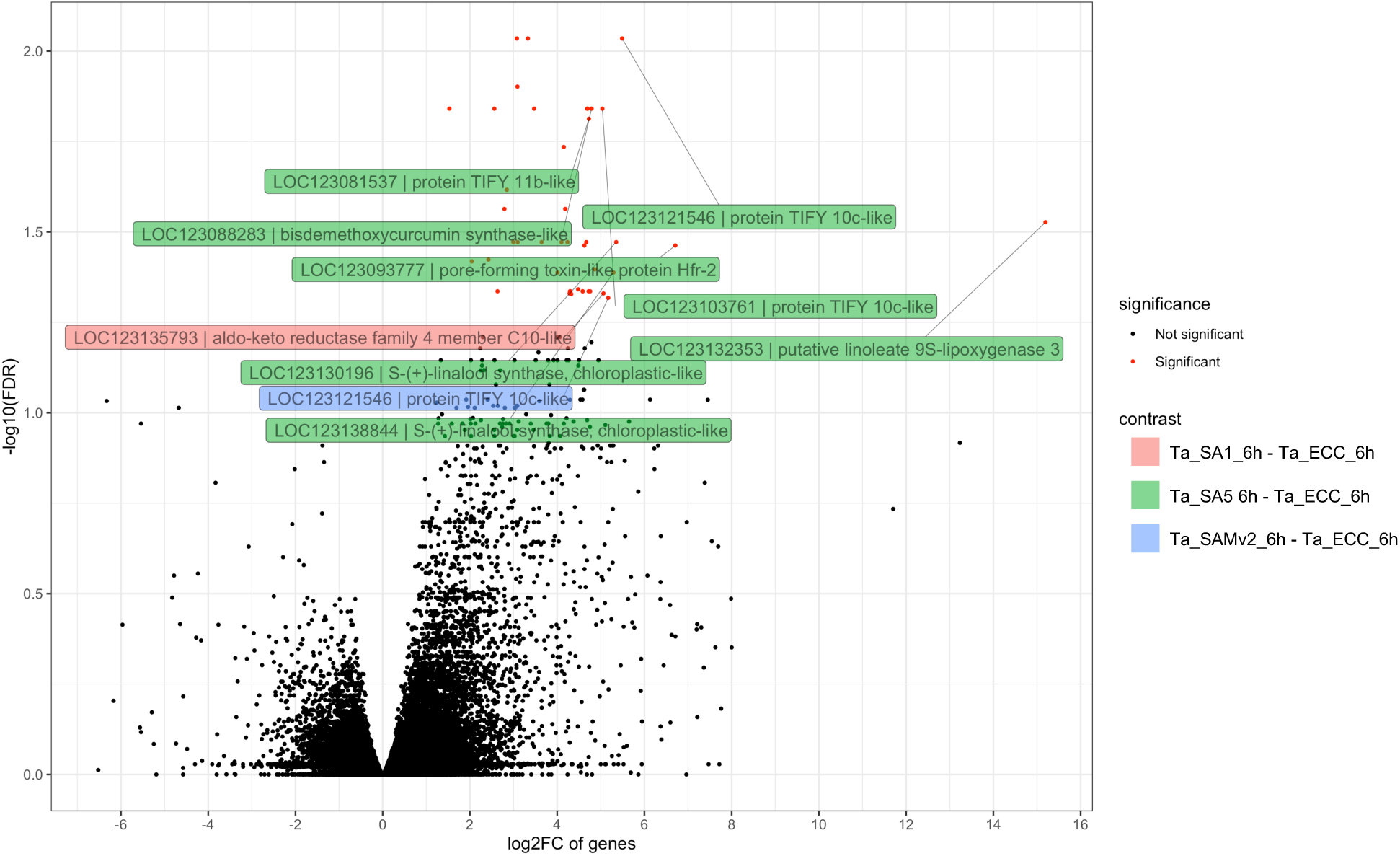
Differentially expressed *Triticum aestivum* genes between the 6 h clip cage control (ECC) and three *Diuraphis noxia* biotypes, SA1, SAMv2 and SA5. Contrast groups include: Ta_SA1_6h - Ta_ECC_6h, Ta_SAMv2 6h - Ta_ECC_6h and Ta_SA5_6h - Ta_ECC_6h. Low expression filtered; adjusted *p*-value < 0.05; log2 fold change > 1; labels: gene ID and product description of top 10 distance to 0, 0.

**Figure 10.**
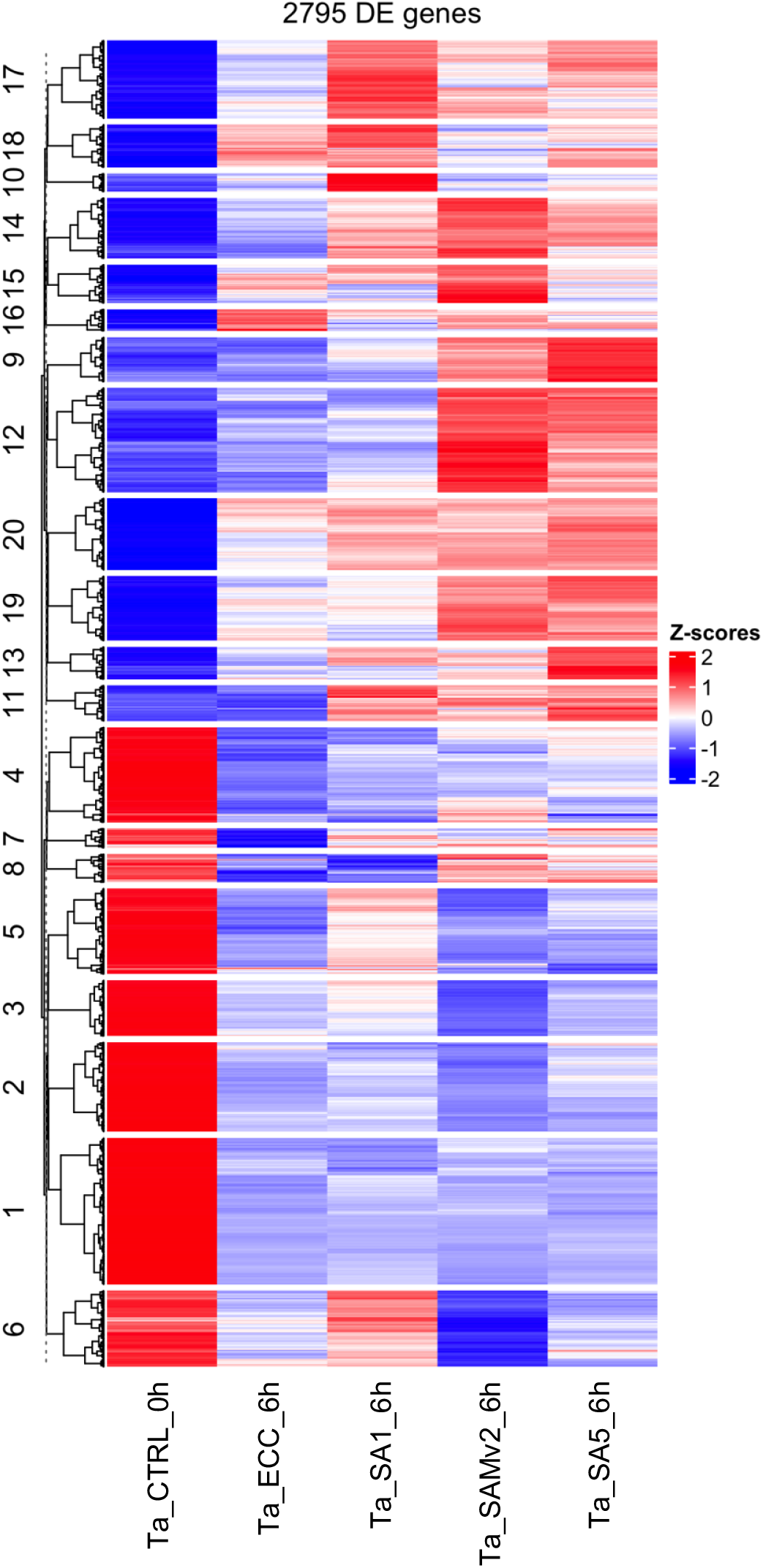
Heatmap demonstrating expression patterns of differentially expressed genes in *Triticum aestivum* cultivar Gamtoos-R*. Diuraphis noxia* biotypes SA1, its direct progenitor SAMv2 and SA5 were contained on the *T. aestivum* leaves for a period of six hours before leaves were sampled. Hierarchical clustering was performed using Euclidean distances measurement and Ward’s method. ECC, empty clip cage control.

Other genes in cluster 11 included 4 paralogues of 1-deoxy-D-xylulose-5-phosphate synthase 2 (DXS2) and S-(+)-linalool synthase. A mitogen-activated protein kinase kinase kinase 18-like (MAP3K18), a aldo-keto reductase family 4 member C10 and the gene encoding for pore-forming toxin protein (HFR-2) are also included in cluster 11 (**Table S10**).

Seven genes DE between SA5 feeding and the ECC were present in cluster 9. DE genes unique to this cluster encode for phenylalanine ammonia-lyase, glutathione S-transferase 6 and the *A. thaliana* MAX1 ortholog, cytochrome P450 711A1 which is present in the strigolactone synthesis pathway (**Table S10**).

In cluster 12 the gene encoding bisdemethoxycurcumin synthase was upregulated after SA5 feeding, but not significantly for the other biotypes. *D. noxia* biotype SA5 induced differential expression of more genes than the other biotypes compared to the 6 h control (**Table 3**). The degree of upregulation was also greater in leaves on which biotype SA5 was contained (**Figure 9**, **Table S10**).

#### *Diuraphis noxia* feeding induced alternative splicing in *Triticum aestivum*

Alternative splicing was calculated as the change in abundance of each transcript compared to the change in gene abundance between conditions. These *p-*values are then combined to a single gene level *p-*value. In contrast to DE, many differential alternative splicing (DAS) events were observed between leaves fed on by different *D. noxia* biotypes. In fact, the inter-biotype contrasts resulted in the largest number of DAS events in *T. aestivum* (**Table 3**, **Table S11**). However, in contrast to DE genes, many DAS genes (49.7% of DAS genes from all contrasts) were uncharacterised.

#### Pathway analysis

Several pathways were found to be significantly enriched for *T. aestivum* genes DE between the 0 h control and the empty clip cage control or *D. noxia* feeding after 6 h. Among others, the KEGG pathways for α-Linolenic acid metabolism (**Figure 11**), terpenoid backbone biosynthesis (**Figure 12**), monoterpenoid biosynthesis (**Figure 13**) and diterpenoid biosynthesis (**Figure 14**) were enriched after SAMv2 and SA5, all biotypes, SA1 or SA5 feeding compared to the 0 h control, respectively (**Table 4**). Plant Reactome pathways enriched within *T. aestivum* after aphid containment included sesquiterpene volatiles biosynthesis, jasmonic acid biosynthesis, abscisic acid mediated signalling and flavin (FMN and FAD) biosynthesis (**Table S12**). Of the contrasts between the 6 h empty clip cage control compared to aphid feeding, only SA5 feeding resulted in the significant enrichment of the Plant Reactome pathway for oleoresin sesquiterpene volatiles biosynthesis (*p*-value < 0.0001) and the KEGG pathways for sesquiterpenoid and triterpenoid biosynthesis (*p*-value < 0.001) and monoterpenoid biosynthesis (*p*-value < 0.05).

**Figure 11.**
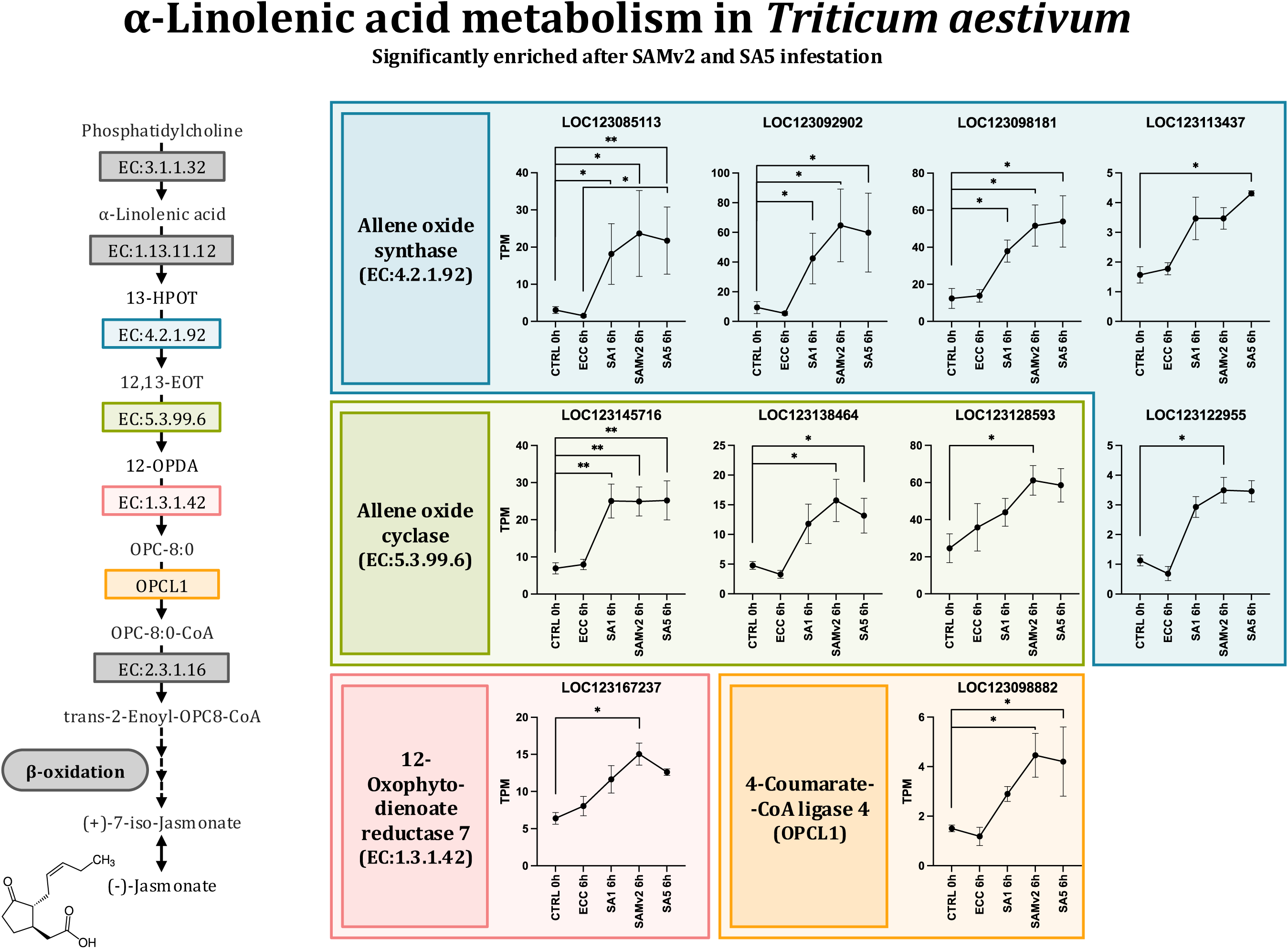
The α-linolenic acid metabolism pathway in *Triticum aestivum* was significantly enriched with differentially expressed genes from the following contrasts: Ta_SAMv2_6h - Ta_CTRL_0h, *p =* 0.0162; Ta_SAM5_6h - Ta_CTRL_0h, *p* = 0.0034.

**Figure 12.**
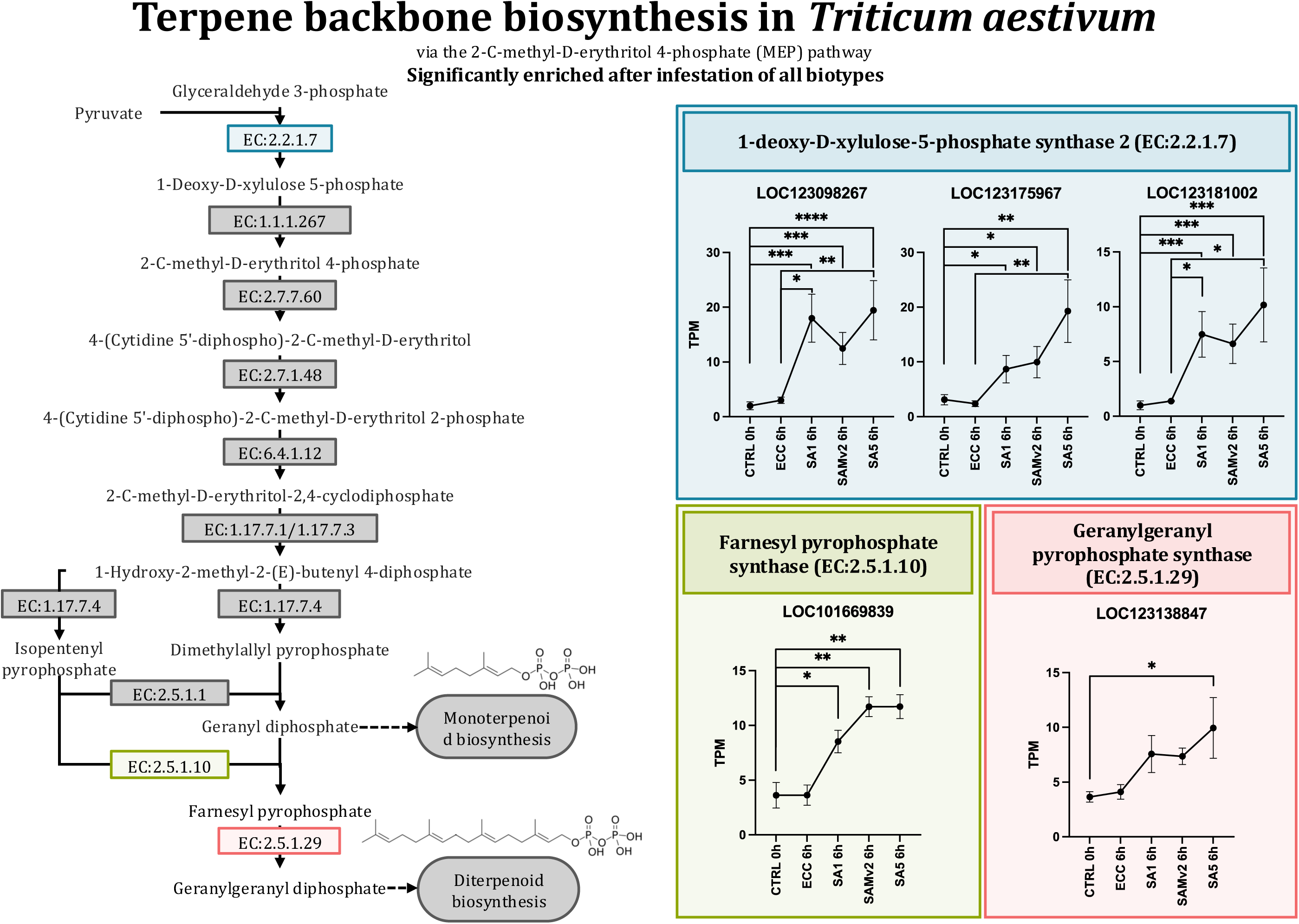
The terpene backbone synthesis (2-C-methyl-D-erythritol 4-phosphate) pathway in *Triticum aestivum* was significantly enriched with differentially expressed genes from the following contrasts: Ta_SA1_6h - Ta_CTRL_0h, *p* = 0.0463; Ta_SAMv2_6h - Ta_CTRL_0h, *p* = 0.0258; Ta_SA5_6h - Ta_CTRL_0h, *p* = 0.0034.

**Figure 13.**
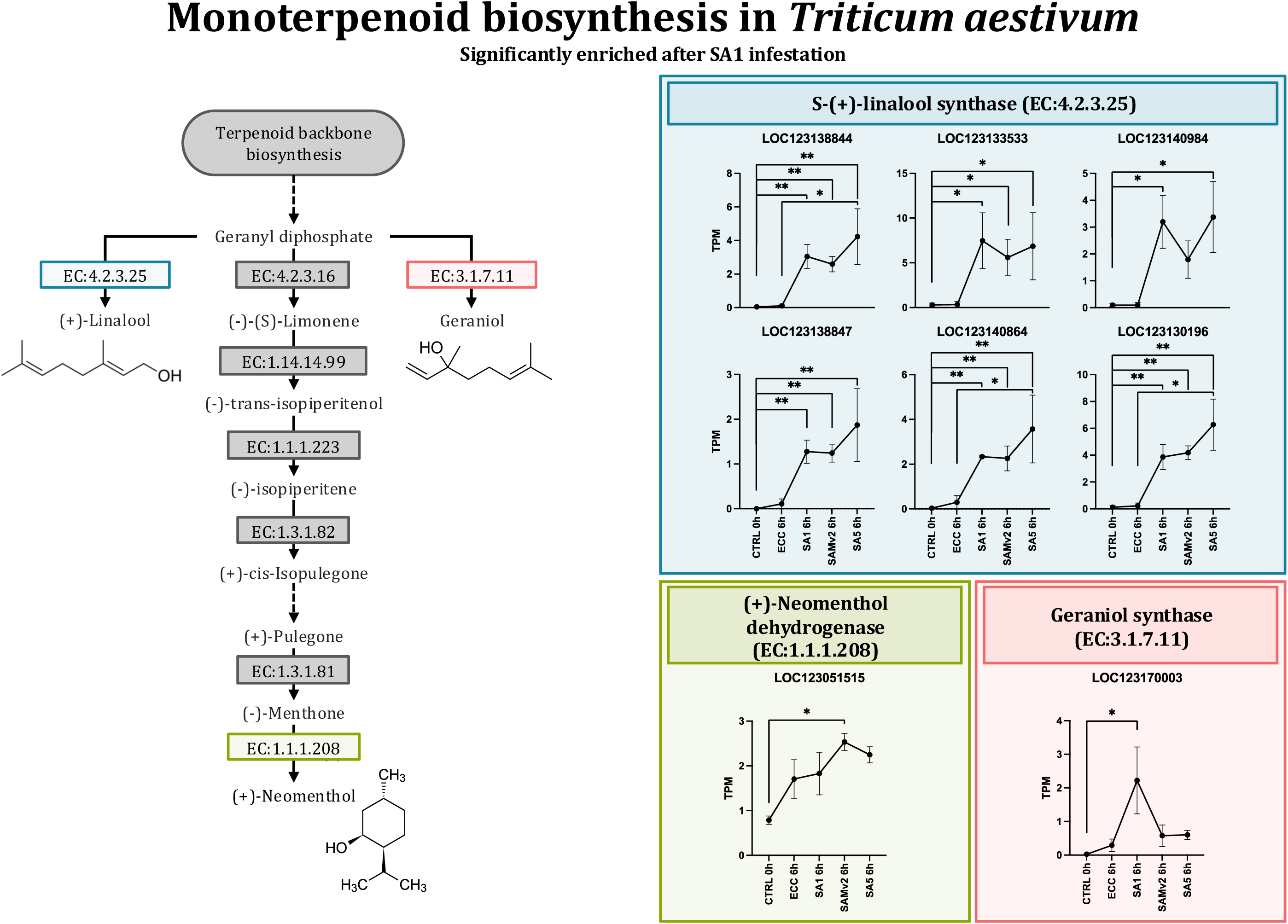
The monoterpenoid biosynthesis pathway in *Triticum aestivum* was significantly enriched with differentially expressed genes from the following contrast: Ta_SA1_6h - Ta_CTRL_0h, *p* = 0.0178.

**Figure 14.**
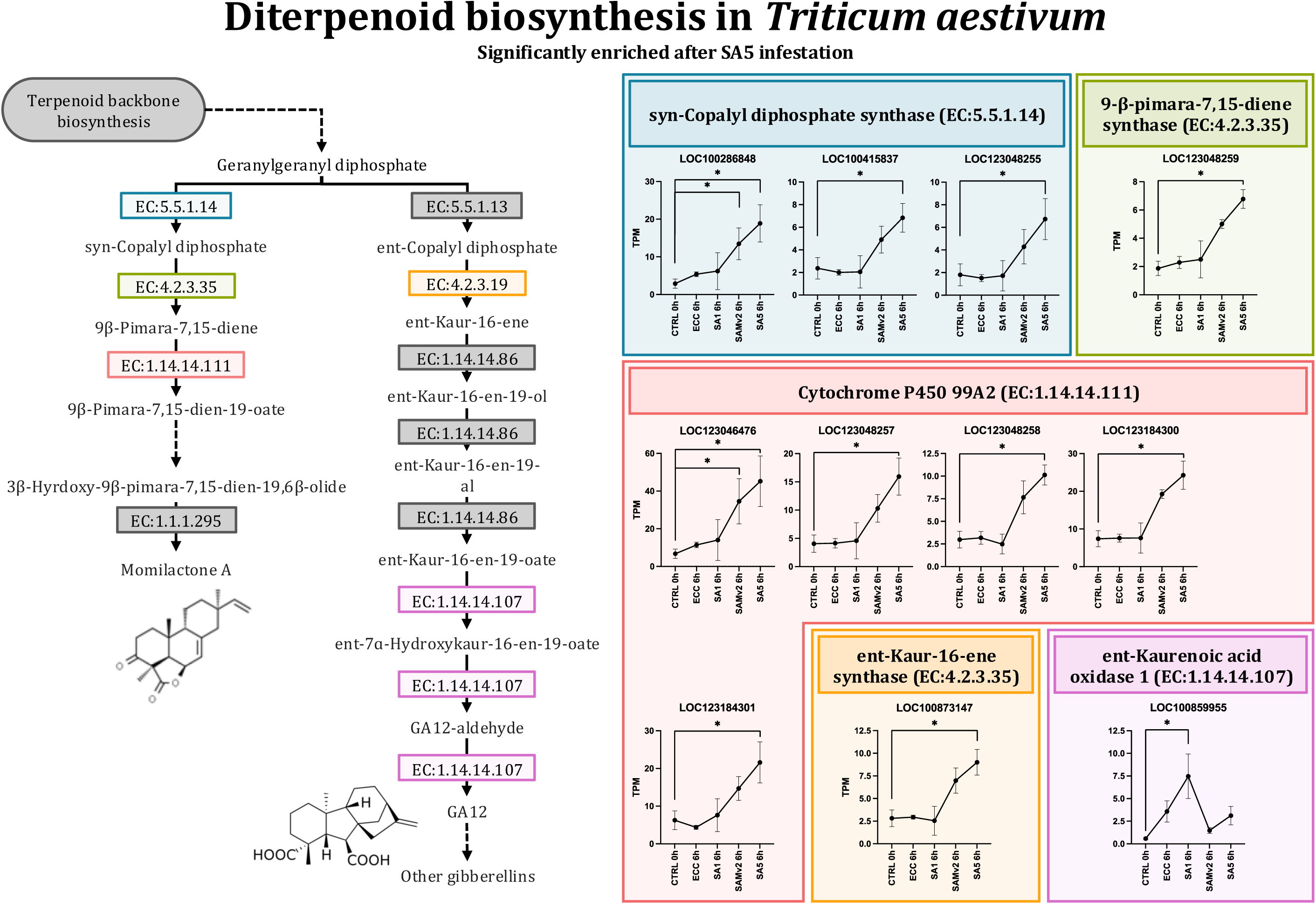
The diterpenoid biosynthesis pathway in *Triticum aestivum* was significantly enriched with differentially expressed genes from the following contrast: Ta_SA5_6h - Ta_CTRL_0h, *p* < 0.00001.

**Table 4.**
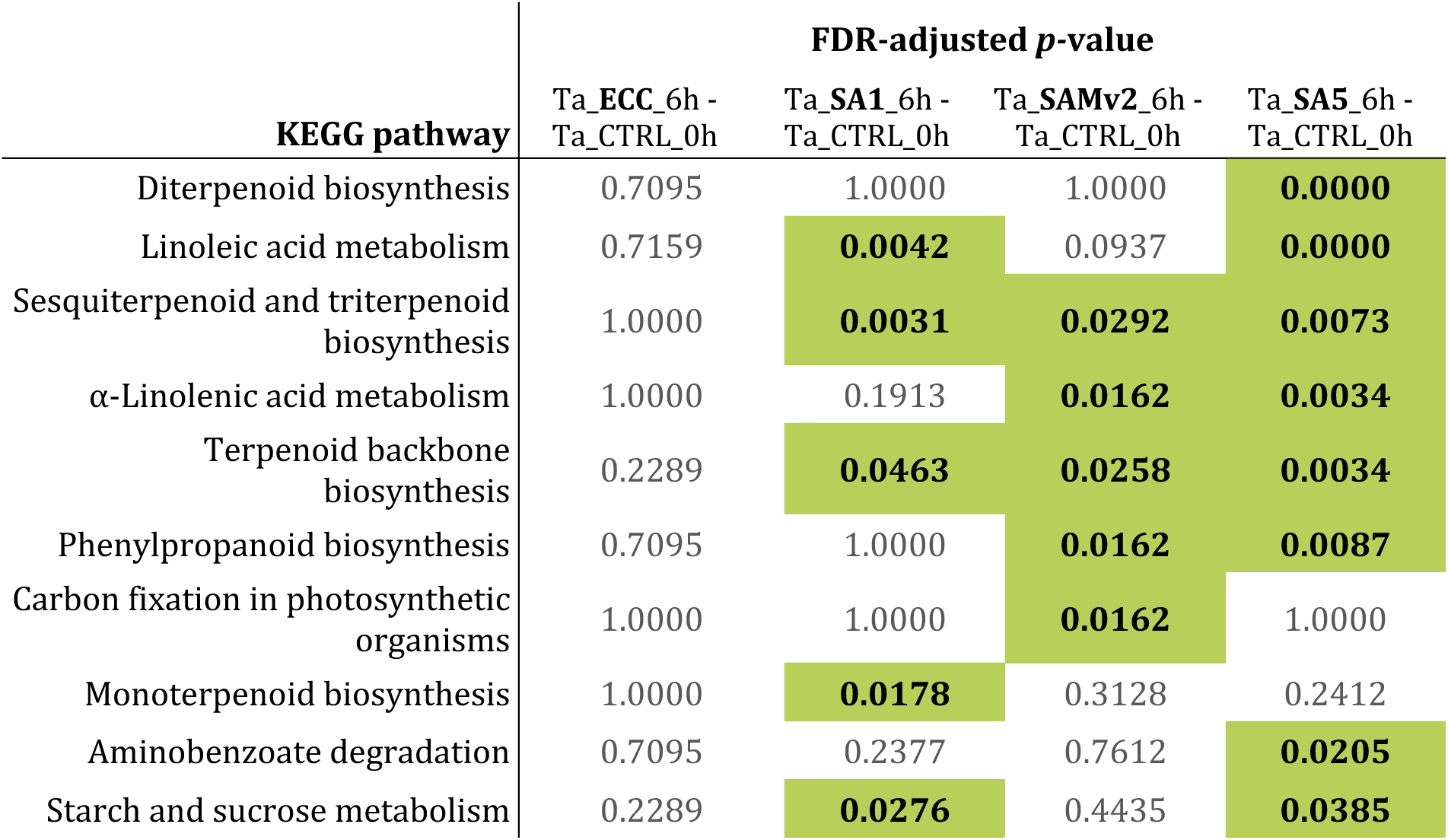
KEGG pathways significantly enriched with *Triticum aestivum* differentially expressed genes from all contrasts between the 0 h control and other conditions (Ta_ECC_6h - Ta_CTRL_0h, Ta_SA1_6h - Ta_CTRL_0h, Ta_SAMv2_6h - Ta_CTRL_0h, Ta_SA5_6h - Ta_CTRL_0h). Enzyme codes and Gene Ontology terms were linked to KEGG pathways. Presented *p-*values were obtained by performing pathway enrichment analyses using Fisher’s test followed by adjustment for FDR. Highlighted *p*-values > 0.05.

### qPCR confirmed RNAseq results

To confirm DE of genes found with RNAseq, qPCR was performed on nine *D. noxia* and ten *T. aestivum* genes to determine relative transcript abundance. Primers for *D. noxia* were designed on the *de novo* assembled transcriptome. All the statistically significant *D. noxia* DE genes found with RNAseq were replicated with qPCR (**Figure 15**).

**Figure 15.**
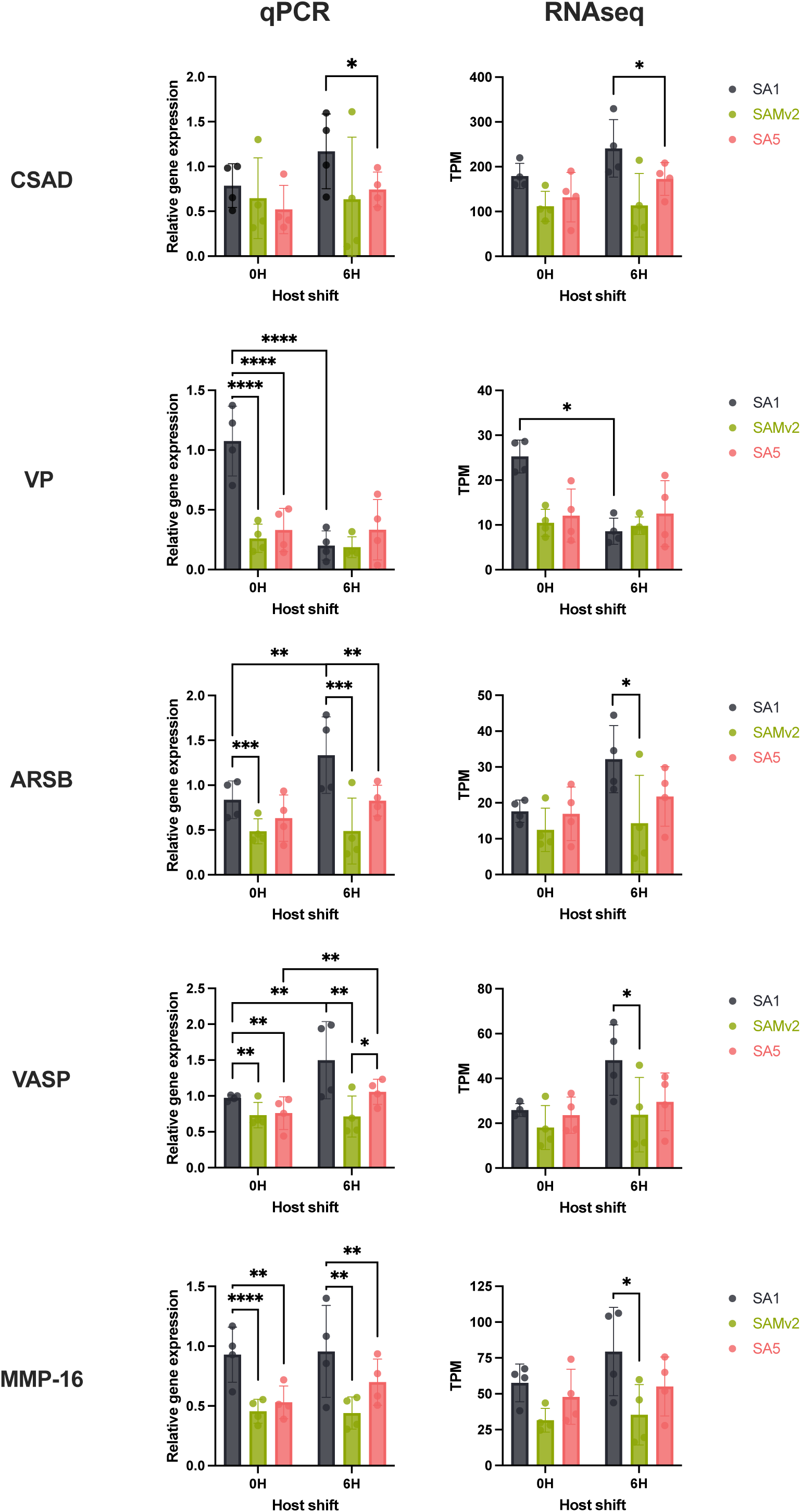
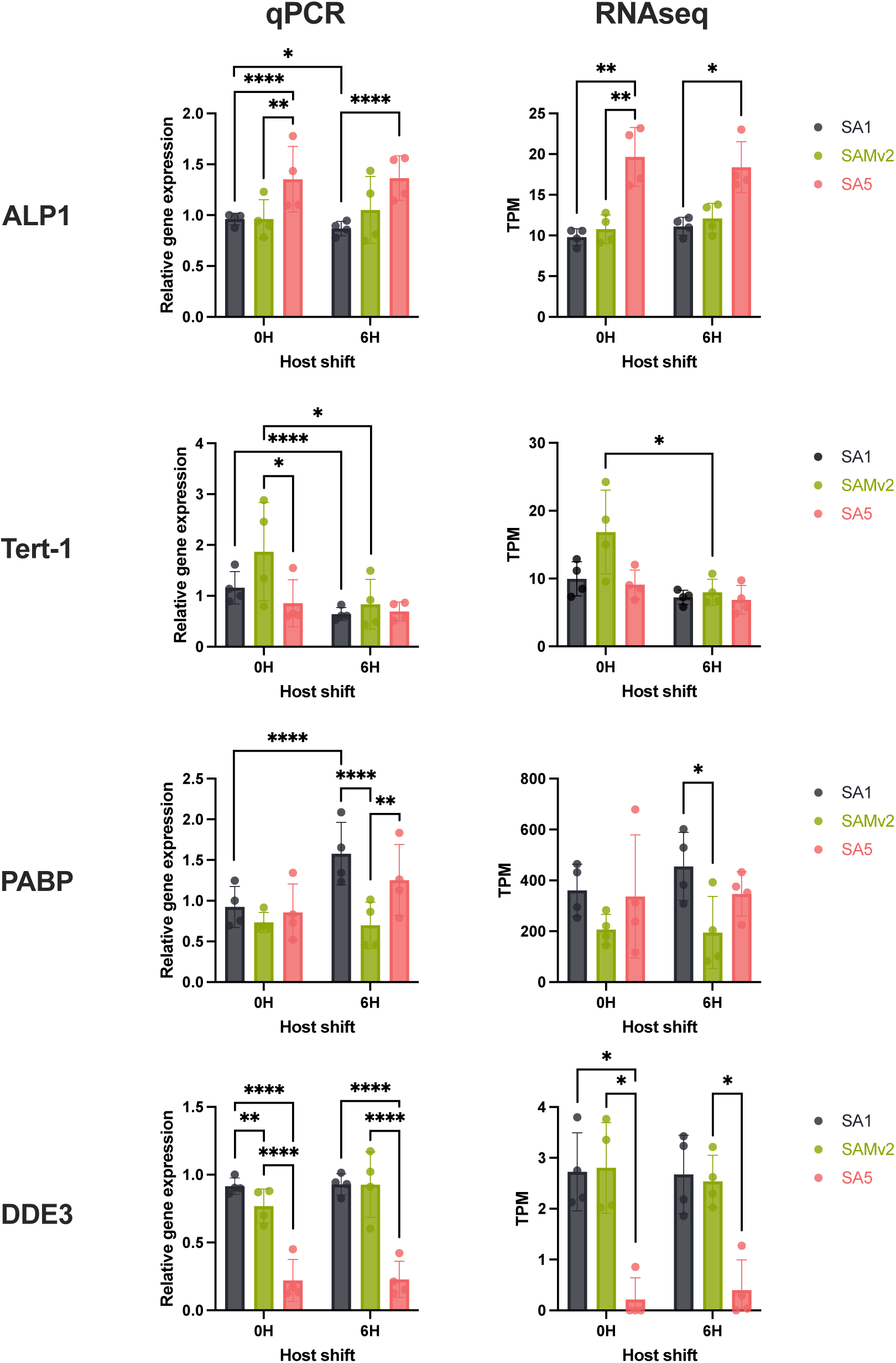
Quantitative PCR confirmed differential expression found with RNAseq of nine *Diuraphis noxia* genes. Primers were designed on the *de novo* assembled *D. noxia* transcriptome. Target gene expression was normalised against the 60S ribosomal proteins L27 and L32 using the Pfaffl method (Pfaffl, 2001). ARSB, arylsulfatase B-like; VP, venom protease-like; MMP-16, matrix metalloproteinase-16; APL-1, protein ALP1-like; DDE3, DDE 3 domain-containing protein; PABP, polyadenylate-binding protein 1-B; Tert-1, facilitated trehalose transporter *Tret1*; VASP, vasodilator-stimulated phosphoprotein; CSAD, cysteine sulfinic acid decarboxylase. Mean ± SD (*n* = 4) Only significant pairwise comparisons shown (using function mcppb20 from WRS2 R package; *, *p* < 0.05; **, *p* ≤ 0.01; ***, *p* ≤ 0.001; ****, *p* ≤ 0.0001).

As the *T. aestivum* Chinese Spring reference transcriptome was used for RNAseq transcript quantification, the corresponding Gamtoos-R transcripts was identified design template-specific qPCR primers. Again, transcript quantification using qPCR corroborated all the significant DE *T. aestivum* genes found with RNAseq analysis (**Figure 16**).

**Figure 16.**
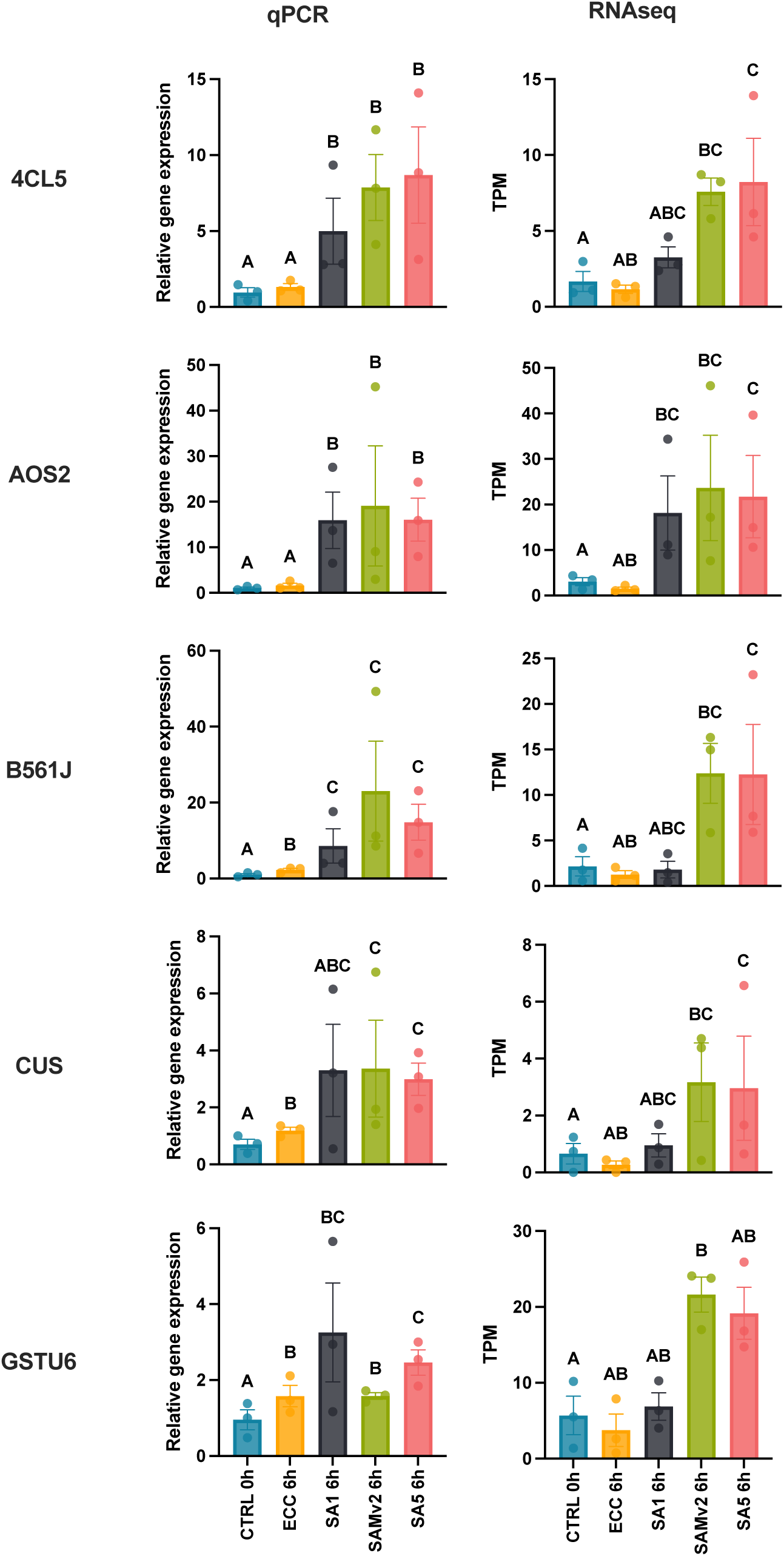
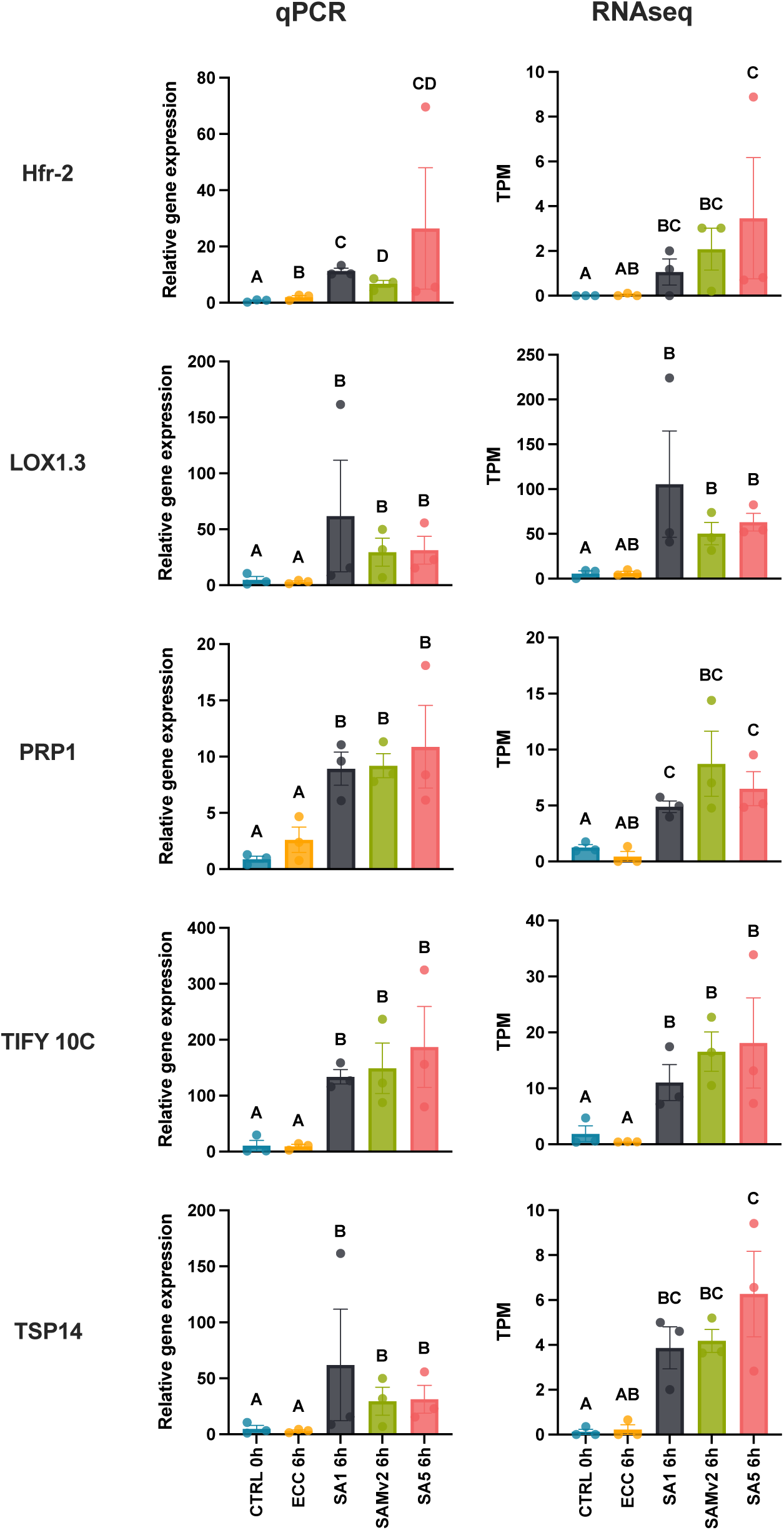
Quantitative PCR was used to confirm differential expression of ten *Triticum aestivum* cultivar Gamtoos-R genes found with RNAseq. The *de novo* assembled Gamtoos-R transcriptome was used for primer design. Target gene expression was normalised against 18S and GAPDH using the Pfaffl method (Pfaffl, 2001). 4CL5, 4-coumarate--CoA ligase 5; AOS2, allene oxide synthase 2; B561J, cytochrome b561 and DOMON domain-containing protein; CUS, bisdemethoxycurcumin synthase; GSTU6, probable glutathione S-transferase *GSTU6*; Hfr-2, pore-forming toxin-like protein *Hfr-2*; LOX1.3, putative linoleate 9S-lipoxygenase 3; PRP1, pathogenesis-related protein 1; TIFY 10C, protein TIFY 10c; TPS14, S-(+)-linalool synthase, chloroplastic. Mean ± SD (*n* = 3) Different letters indicate significant pairwise comparisons (using function mcppb20 from WRS2 R package, *p* < 0.05).

## Discussion

To better understand the molecular interaction between *D. noxia* and *T. aestivum* with specific focus on the formation of virulence in *D. noxia* to resistant *T. aestivum* cultivars, an RNAseq analysis was performed on virulent and avirulent *D. noxia* biotypes before and after transfer from a susceptible to a near-isogenic, *Dn7*-containing *T. aestivum* cultivar. In addition, an RNAseq analysis was also performed on the resistant *T. aestivum* cultivar to which the same aphid biotypes were transferred.

### Diuraphis noxia

To study the molecular basis of virulence in *D. noxia* to resistant *T. aestivum*, we compared the transcriptional profiles of the virulent SAMv2 biotype and its progenitor, SA1. Since SAMv2 is a descendant of SA1, we expected to find few differences between them, with most of the observed changes being related to the development of virulence in SAMv2. In addition, the stress of transferring the biotypes from susceptible to resistant host could elicit differential transcriptional regulation which in turn should make the inter-biotype differences more pronounced. Indeed, no significant DE unigenes were found when comparing SAMv2 to its progenitor SA1 after feeding on Gamtoos-S. In contrast, 19 unigenes were DE after transfer to Gamtoos-R. All of these were upregulated in SA1 relative to SAMv2. The predicted functions these unigenes indicates this it is involved in response to stress. One such gene, haemolymph juvenile hormone binding (HJHB), transports insect juvenile hormone (JH) to its active site and protects it from degradation (Hidayat & Goodman, 1994). In *Drosophila*, JH inhibits both metamorphosis in juveniles as well as embryogenic development (Riddiford & Williams, 1967) while allowing growth and moulting. Elevated JH has been associated with the continuation of parthenogenetic morph production in *Acyrthosiphon pisum*, while a decrease in JH results in the production sexual morphs (Ishikawa *et al*., 2012). Ishikawa et al. (2013) found that a decrease in JH resulted in the development of winged morphs in *Megoura crassicauda*. However, wing formation in *D. noxia* like other aphids are typically associated with high population densities, not feeding on resistant host plants. Furthermore, a male *D. noxia* has not yet been observed and thus if it does produce sexual nymphs, it must be a rare event. Thus, if HJHB does result in elevated levels of JH as proposed by (Hidayat & Goodman, 1994), it may function to delay embryonic development in *D. noxia* SA1 when feeding on a resistant host. Upregulation of *CYP4C1* (cytochrome P450 4C1) in SA1 compared to SAMv2 post host shift was also observed. Differential expression of *CYP4C1* is associated with *Sitobion avenae* (English grain aphid) feeding on *T. aestivum* compared to *Hordeum vulgare* (Huang *et al*., 2019) and insecticide resistance in other insects (Kim Lien *et al*., 2019; Bouafoura *et al*., 2022).

It should be considered that the differential response of a virulent aphid when feeding on a resistant host may simply be owing to it being able to feed unhindered and not the underlying reason for it being able to feed more effectively. Thus, the contrast discussed above (DnSAMv2_6h - DnSA1_6h) include stress responses by SA1 owing to its reduced ability to feed on *D. noxia* resistant *T. aestivum*. This does not help us understand how SAMv2 is adapted to feed on *D. noxia* resistant *T. aestivum* — a question better answered by comparing the average TPM of 0 h and 6 h between SA1 and SAMv2 in the following contrast: (DnSAMv2_0h + DnSAMv2_6h)/2 - (DnSA1_0h + DnSA1_6h)/2. Here, more stable gene expression differences between SA1 and SAMv2 were found with less association to the host shift and thus SA1’s response to its reduced ability to feed. DE unigenes from this contrast include *L-xylulose reductase*, upregulated 3.5-fold in SAMv2. L-xylulose reductase catalyses the reduction of L-xylulose to xylitol, but importantly can also utilise other substrates. It was found to be upregulated in virulent *Schizaphis graminum* (greenbug) relative to avirulent biotypes (Pinheiro *et al*., 2014) and DE between different *Rhopalosiphum maidis* (corn aphid) linages feeding on *Zea mays* (Guo *et al*., 2022). MacWilliams et al. (2020) identified L-xylulose reductase as an effector from *Aphis craccivora* (cowpea aphid) saliva. It was shown *in vitro* to not only reduce xylulose, but also methylglyoxal which accumulates in *Vigna unguiculata* (cowpea) during *A. craccivora* feeding and is cytotoxic at elevated levels.

Six of the seven DE unigenes in cluster 7 are of unknown function and four are not predicted to be protein-coding. Transcript GG_757_c169_g1_i1 is predicted to produce a 426 aa protein of unknown function. Interestingly, it is expressed 17.4-fold higher in SAMv2 relative to SA1 (the greatest difference in DE between SAMv2 and SA1), is fairly abundant at 36.8 TPM (ranked within the top 89% of transcripts expressed in SAMv2) and is potentially secreted as it contains a signal peptide. Further investigation into the function of this transcript is recommended.

The transfer of *D. noxia* from the susceptible host Gamtoos-S to a near-isogenic line containing the *Dn7* resistance gene resulted in the DE of 11 unigenes with diverse functions. These grouped in clusters 9 and 4. As Solute carrier family 26 member 10-like, an anion transporter, was downregulated in all biotypes, 2.2-fold in SAMv2 after the host shift. *Tert-1* (encoding for facilitated trehalose transporter) was also downregulated in SAMv2 after the host shift. The DE of aforementioned membrane transporters 6 hours after the host shift may be explained by the interruption of phloem feeding and/or different sugar and amino acid concentrations in the resistant cultivar. Before the phloem sieve element is found, aphids often probe other tissues like the xylem or mesophyll which have different sugar and mineral contents. Tert-1 facilitates trehalose transport from the insect fat body (where trehalose is synthesised) to the haemolymph and thus regulates haemolymph trehalose concentration (Kanamori *et al*., 2010). Trehalose is the main sugar present in insect haemolymph and functions as an energy store, stabilises proteins during osmotic or thermal stress and regulates feeding behaviour (Thompson, 2003). *Tert-1* was also found to be differentially expressed in *S. avenae* feeding on resistant compared to susceptible *T. aestivum* cultivars (Lan *et al*., 2021) and multiple *Tert-1* paralogues were differentially regulated in *M. persicae* feeding on *Brassica rapa* compared to *Nicotiana benthamiana* (Mathers *et al*., 2017).

Transposases were also differentially regulated 6h after the host shift including a P-element transposase and a DDE domain-containing transposase. DE of TEs is expected when comparing between aphid biotypes as aphids mostly reproduce asexually and thus require other mechanisms to introduce genetic variation (Singh *et al*., 2020; Panini *et al*., 2021). Indeed, DE of TEs were found here in all contrasts tested, with the highest number of DE events present when comparing SA5 to SA1 or SAMv2. Baril et al. (2023) recently found that gene families associated with the detoxification of xenobiotics (synthetic insecticides and toxic plant metabolites) were enriched with TEs. Indicating its importance in the evolutionary adaption of aphids to xenobiotic metabolism. It is however interesting that within 6 h of the host shift the unigenes with the highest degree of DE encoded for transposons. P-element transposase was upregulated 12.6-fold while the DDE domain-containing transposase was downregulated 17.8-fold. This may indicate that transposons are responsible for short-term gene regulation through transcriptional activation and not only genetic adaptation. It should be noted that contrasts discussed here, DnSA1_6h - DnSA1_0h; DnSAMv2_6h - DnSAMv2_0h; (DnSA1_6h + DnSAMv2_6h)/2 - (DnSA1_0h + DnSAMv2_0h)/2, does not allow for one to distinguish between the effect of transferring to a resistant cultivar from simply transferring the aphids to a different plant of the same (susceptible) cultivar.

Comparison of SA5 to SA1 or SAMv2 resulted in more transcriptional regulation events than SA1 compared to SAMv2. This is expected as SA5 is more distantly related to SA1 and its descendent, SAMv2. Unigenes DE between SA5 and the other biotypes mainly encoded for transposons (7 of 17 DE unigenes) as well as histone acetyltransferase, the ortholog for *Drosophila* ring canal kelch protein (*kel*) and uncharacterised proteins. *kel* is involved in the formation of intercellular ring canals connecting oocytes and the surrounding nurse cells. Through the ring canals, cytoplasmic components are provided to the oocytes from the nurse cells (Xue & Cooley, 1993). The absence of *kel* results in female sterility (Robinson & Cooley, 1997). The function of this *D. noxia* kelch protein and the effect its increased expression in SA5, will need to be confirmed experimentally.

### Triticum aestivum

Clip cages are used in many entomological studies especially when the interaction between host plant and insect is investigated. Here, it was found that differential gene and transcript expression in *T. aestivum* resulting from placing an empty clip cage on its leaf was induced to a similar amount, and most genes (54%) shared identity with genes DE when *D. noxia* was contained on its leaf using clip cages. The DE of these shared genes will thus be more difficult to associate with insect feeding alone when the 6 h ECC is compared with 6 h aphid feeding. The physiological effect of the clip cage on the plant may also influence its response to aphid feeding, as it is known that JA signalling is induced during wounding (by the clip cage) and also acts antagonistically to SA signalling. Thus, the use of clip cages should be done with caution and the effect of its use on the plant should be accounted for in the experimental design.

Even though the clip cage influenced *T. aestivum* gene regulation, pathways distinct to *D. noxia* feeding were enriched with DE genes between 0 h control and the other treatments. The pathways for alpha-linolenic acid metabolism was enriched in leaves fed on by SAMv2 and SA5 which results in the biosynthesis of JA (specifically enriched in SAMv2). Seventeen DE genes involved in jasmonic acid synthesis or signalling clustered together (cluster 11 on the heatmap of *T. aestivum* DE genes) and included the TIFY proteins 10C, 10B and 11B, transcription factors bHLH 6 and 148, and allene oxide synthase 2 (AOS2). TIFY proteins in *O. sativa*, specifically 10C, acts as a repressor of transcription factors in the absence of JA. In the presence of JA, these proteins are degraded which leads to rapid activation of JA-responsive genes such as those encoding S-(+)-linalool synthase (TPS14). Upregulation of TIFY 10C therefore acts to regulate the JA-induced defence response (Yamada *et al*., 2012; Taniguchi *et al*., 2014). JA regulates the production of many secondary metabolites including terpenes (Avanci *et al*., 2010).

Pathways for the biosynthesis of the terpene backbone, monoterpenes, diterpenes and sesquiterpenes were upregulated in all *D. noxia* biotypes, SA1, SA5 and all biotypes, respectively. Three paralogues of 1-deoxy-D-xylulose-5-phosphate synthase 2 (DXS2) were DE in response to aphid feeding with the largest fold-change in expression seen for SA5. DXS2 catalyses the first and rate-limiting reaction in the 2-*C*-methyl-D-erythritol-4-phosphate pathway which results in the synthesis of the terpene backbone (Tian *et al*., 2022).

Six genes encoding TPS14 from the monoterpenoid pathway were upregulated in *T. aestivum* when fed on by all biotypes. Increased S-(+)-linalool in an octoploid Trititrigia was correlated with reduced *S. avenae* fecundity and the monoterpenoid also had a significant repellent effect on the aphid (Zhan *et al*., 2021). This supports earlier phenotypic findings that *Dn7* resistance is based on antibiosis (Lazzari *et al*., 2009) and antixenosis (Anderson *et al*., 2003; Haley *et al*., 2004). In addition, a gene encoding (+)-neomenthol dehydrogenase was upregulated in *T. aestivum* leaves fed on by SAMv2. This enzyme produces the monterpene alcohol, (+)- neomenthol, the effect of which on aphids has not been studied as a single component, but as a component of *Mentha × piperita* essential oil it has been observed to deter *A. pisum* (Lacotte *et al*., 2023), *M. persicae* (Dancewicz *et al*., 2021), *Macrosiphum euphorbiae* (Cantó-Tejero *et al*., 2022a) and *Rhopalosiphum padi* (Pascual-Villalobos *et al*., 2017) or be toxic to *Aphis gossypii* (Ebadollahi *et al*., 2017; Heydari *et al*., 2020). The abundance of (+)-neomenthol in *T. aestivum* infested with aphids or the effect thereof on *D. noxia* has not been investigated.

The gene encoding geraniol synthase was upregulated 58-fold in leaves fed on by SA1 relative to the 0 h control. An *Oryza sativa* geraniol synthase homolog was found to be upregulated in response to JA. Additionally, geraniol produced by geraniol synthase, had a significant negative effect on the growth of *Xanthomonas oryzae*, the causal agent of rice blight disease (Kiyama *et al*., 2021). The production of geraniol in *T. aestivum* in response to JA or aphid feeding has not been demonstrated. However, toxicity of topically applied geraniol against *M. persicae* has also been shown: 0.6% v/v geraniol resulted in 52.5% lethality (Cantó-Tejero *et al*., 2022b).

The diterpenoid synthesis pathway was significantly enriched in *T. aestivum* leaves fed on by SA5. Specifically, genes encoding enzymes in pathways leading to the production of momilactone A and gibberellins were upregulated. Induction of momilactone A accumulation occurs in an herbivore-specific manner (Yactayo-Chang *et al*., 2020) and was specifically found to be induced by the phloem-feeding *Sogatella furcifera* in *O. sativa* (Kanno *et al*., 2012). It functions to deter herbivore feeding and can be toxic to pathogens. Momilactone A accumulation in distinct transgenic *O. sativa* lines were shown to increase resistance to both *Magnaporthe grisea* (rice blast fungus) and *X. oryzae* (Sawada *et al*., 2004; Mori *et al*., 2007).

Following feeding of SA5, the *A. thaliana max1* ortholog *CYP711A1* was upregulated 3-fold relative to the ECC sample. *CYP711A1* is responsible for the synthesis of canonical and non-canonical strigolactones by converting carlactone into carlactonoic acid (Zhang *et al*., 2014; Abe *et al*., 2014). Other genes DE relative to those DE from the ECC sample, which were also present in cluster 9 are all directly involved in biotic stress. For example, phenylalanine ammonia-lyase is involved in SA synthesis while 4-coumarate-CoA ligase 5 partakes in the synthesis of JA. Cross-talk between SA and JA with Strigolactone has been speculated, but the mechanisms are not known (Qi *et al*., 2024). This may be the first report of aphid feeding-induced upregulation of *CYP711A1* that points to the involvement strigolactones in biotic stress.

## Acknowledgements

This research was supported by a Commonwealth Split-site Scholarship (ZACN-2018-353) funded by the UK government and a DST-NRF Innovation Doctoral Scholarship (118101) funded by the National Research Foundation of South Africa.

## Author contributions

HWS designed and performed all experimental work, analysed and interpreted all results, discussed data and wrote the manuscript. CHF critically reviewed the manuscript. AMB conceptualised the idea and critically reviewed the manuscript.

## Data availability

The RAW RNAseq data were deposited at the BioProject database: accession number PRJNA1120216.

The assembled *D. noxia* and *T. aestivum* cultivar Gamtoos-R transcriptomes are also available at the above BioProject. The specific TSA the accession numbers are: GKVM01000000 and GKWO01000000.

