## Supplementary Tables for "Investigating the *Diuraphis noxia*-*Triticum aestivum* interaction using transcriptomics"

**Table S1**. Primers used for qPCR to confirm significant differentially expressed *Triticum aestivum* genes found through RNAseq.

| IWGSC transcript | Gamtoos-R transcript | Name | Forward primer | Reverse primer | Product Size | $T_a$ |
|---|---|---|---|---|---|---|
| XM_044474884 | GG_83993_c0_g1_i1 | 4CL5 | ATGGTGCTTGATTATTTGTGTGAGA | GGCGACTGATTGATCCTTGC | 110 bp | 60°C |
| XM_044506705 | GG_97134_c0_g1_i1 | AOS2 | GTAGTAGTACGAGTCAGTCATGCA | AACACACACACATACACATACACAC | 110 bp | 60°C |
| XM_044520398 | GG_104417_c0_g1_i1 | B561J | CCAACTTGCAGAGCCTTATGAG | GGATGACACTAACGGATAATTACACC | 120 bp | 60°C |
| XM_044601233 | GG_77805_c0_g1_i1 | BHLH6 | AGTCTCTCTGTTGCAAGGCTAA | TGCAGATTCACTCATTTTGCTCC | 110 bp | 60°C |
| XM_044510480 | GG_99629_c0_g1_i3 | CUS | GATCTTTGTGCTCGATGAGGTC | CAGTGAACCCCGGTCCAA | 110 bp | 60°C |
| XM_044505259 | GG_99521_c0_g1_i1 | CYP711A1 | GTCAAGCTCCAGGTCGTCC | ATCGATGTTCAACAGTTCAAGTAGC | 109 bp | 60°C |
| XM_044520220 | GG_5401_c0_g1_i1 | DXS2 | ATCCTCATCACCGCCGAG | CGGTAGAAACATCGATCTCAGC | 117 bp | 60°C |
| XM_044593668 | GG_74677_c0_g1_i1 | GSTU6 | CTGCAACCACTTTCATCAACGTC | GTCCACACGCCCAGCAG | 110 bp | 60°C |
| XM_044515830 | GG_34636_c0_g1_i2 | Hfr-2 | GCCAAGTTAAGAGCCGGGAT | GCCTCTTGTGATGTCGTTGTC | 116 bp | 60°C |
| XM_044552125 | GG_115683_c0_g1_i2 | LOX1.3 | ACCTCGGCAAGCTCAATGTT | TGTTTGGGAGGCTGTTGATCT | 111 bp | 60°C |
| XM_044469391 | GG_14346_c0_g1_i4 | PAL | CTAACATCCTTAGCCTCCTTGTC | CGGGGTGGTGCTTCAATTT | 111 bp | 60°C |
| XM_044472362 | GG_17520_c0_g1_i1 | PRP1 | ACTATCAAAGGGTGTCATGCTGTT | TAAAGCAAACAAATATTTCCTTTTATTCCGG | 102 bp | 60°C |
| XM_044541547 | GG_45574_c0_g1_i2 | TIFY 10C | TCGTCGTAGGAGAAGGAGCTTT | TGGCCAACACTAGATTTCTCGA | 115 bp | 60°C |
| XM_044550149 | GG_115293_c0_g1_i9 | TPS14 | TCCTTCCCCTTCTCAGTTCAG | CGGAAGAAGGCACGGTTG | 92 bp | 60°C |
| XR_006488544 | GG_15_c0_g1_i7 | 18S | CGTCCCTGCCCTTTGTACAT | AACACTTCACCGGACCATTCA | 63 bp | 60°C |
| XM_044580916 | GG_128368_c0_g1_i2 | GAPDH | GCCAGTTACCGTCTTTGGCGTC | GGCCTTGTCCTTGTCAGTGAAG | 109 bp | 57°C |

**Table S2**. Primers used for qPCR to confirm significant differentially expressed *Diuraphis noxia* genes found through RNAseq.

| **Target ID** | **Name** | **Product description** | **Forward primer** | **Reverse Primer** | **Product size** | **$T_a$** |
|---|---|---|---|---|---|---|
| GG_403_c15_g1 | ARSB | Arylsulfatase B-like | CGAGGCGCTCAAGTACTACA | GTCGGTCAGTACGTCCAACT | 117 bp | 55°C |
| GG_1409_c4_g1 | VP | Venom protease-like | TCAACACGACAAATGGAGAGGAT | CGAGCCACATTCCCAGAACT | 106 bp | 55°C |
| GG_617_c153_g1 | L27 | 60S ribosomal protein L27 | ACCAGCACGATTTTACCAGATTTC | CGTAGCCTGCCCTCGTGTA | 89 bp | 56.3°C |
| GG_757_c6_g1 | L32 | 60S ribosomal protein L32 | CGTCTTCGGACTCTGTTGTCAA | CAAAGTGATCGTTATGACAAACTCAA | 74 bp | 56.3°C |
| GG_1563_c7_g1 | MMP-16 | Matrix metalloproteinase-16 | GGGGCGGGAATGGAAAGATT | GTTGCTGATAGGCTTCGGGT | 107 bp | 58°C |
| GG_713_c29_g1 | APL-1 | Protein ALP1-like | GCCCACAAAGCAAAGGAGAA | CGTTAAATTCACCGACACCGT | 111 bp | 60°C |
| GG_130_c65_g1 | DDE3 | DDE 3 domain-containing protein | ATGCTGGCGACTGTACTACG | CGTTTTCCTTTGCCCGATGG | 115 bp | 60°C |
| GG_1406_c61_g1 | PABP | Polyadenylate-binding protein 1-B-like | ATAAAATGGCGTGTTCATGTCTTG | CGTGTCGAAGTCTCCTAGATCTA | 100 bp | 60°C |
| GG_1562_c83_g1 | Tert-1 | Facilitated trehalose transporter Tret1-like | GTCGTCAAGCCGTATTCCCA | TTGAGGCGATTCGTGCTCAT | 112 bp | 60°C |
| GG_1071_c16_g1 | VASP | Vasodilator-stimulated phosphoprotein-like | CACGATTGTCACTGGATTGCC | GGCTTGTTGTCTCGGATGGA | 102 bp | 61°C |
| GG_1449_c28_g1 | CSAD | Cysteine sulfinic acid decarboxylase | TGATGATCACGTACCAGCCAG | ACCCAGCCGTTCGATTTCAT | 119 bp | 63°C |

**Table S3**. Quality control of *Diuraphis noxia* samples used for RNA sequencing. RNA integrity was confirmed with the 2200 TapeStation System (Agilent). Raw paired-end reads of 150 bp were trimmed for quality and to remove sequencing adapters. Reads below 36 bp were discarded.

| **Treatment** | **Biological replicate** | **RIN value** | **# paired-end reads post trim (M)** | **% paired-end reads retained post trim** | **Avg. quality per read post trim** | **Q30 post trim** |
|---|---|---|---|---|---|---|
| DnSA1_0h | 1 | 9.8 | 14.40 | 99.8% | 35.44 | 98.4% |
| DnSA1_0h | 2 | 9.4 | 31.07 | 99.5% | 35.25 | 97.6% |
| DnSA1_0h | 3 | 9.8 | 16.13 | 99.7% | 35.33 | 97.6% |
| DnSA1_0h | 4 | 9.4 | 10.95 | 99.8% | 35.44 | 98.2% |
| DnSA1_6h | 1 | 9.5 | 11.69 | 99.8% | 35.46 | 98.4% |
| DnSA1_6h | 2 | 9.4 | 31.07 | 99.6% | 35.26 | 97.7% |
| DnSA1_6h | 3 | 9.8 | 12.46 | 99.6% | 35.34 | 97.6% |
| DnSA1_6h | 4 | 9.8 | 10.52 | 99.7% | 35.36 | 98.2% |
| DnSA5_0h | 1 | 8.9 | 16.29 | 99.6% | 35.37 | 97.7% |
| DnSA5_0h | 2 | 9.9 | 10.19 | 99.2% | 35.33 | 98.0% |
| DnSA5_0h | 3 | 9.5 | 10.46 | 99.6% | 35.4 | 98.2% |
| DnSA5_0h | 4 | 9.9 | 31.7 | 99.6% | 35.26 | 97.7% |
| DnSA5_6h | 1 | 9.8 | 14.63 | 99.6% | 35.34 | 97.6% |
| DnSA5_6h | 2 | 9.9 | 10.44 | 99.3% | 35.32 | 97.9% |
| DnSA5_6h | 3 | 9.7 | 10.29 | 99.7% | 35.39 | 98.2% |
| DnSA5_6h | 4 | 9.9 | 31.46 | 99.6% | 35.28 | 97.7% |
| DnSAMv2_0h | 1 | 9.9 | 10.89 | 99.7% | 35.4 | 98.3% |
| DnSAMv2_0h | 2 | 10 | 12.75 | 99.7% | 35.32 | 97.6% |
| DnSAMv2_0h | 3 | 9.4 | 10.22 | 99.7% | 35.39 | 98.2% |
| DnSAMv2_0h | 4 | 8.3 | 30.86 | 99.3% | 35.16 | 97.3% |
| DnSAMv2_6h | 1 | 9.8 | 16.17 | 99.9% | 35.42 | 98.3% |
| DnSAMv2_6h | 2 | 9.4 | 13.88 | 99.6% | 35.34 | 97.6% |
| DnSAMv2_6h | 3 | 9.9 | 10.41 | 99.5% | 35.43 | 98.2% |
| DnSAMv2_6h | 4 | 9.8 | 28.28 | 99.4% | 35.28 | 97.6% |

**Table S4**. Quality control of *Triticum aestivum* samples used for RNA sequencing. RNA integrity was confirmed with the 2200 TapeStation System (Agilent). Raw paired-end reads of 150 bp were trimmed for quality and to remove sequencing adapters. Reads below 36 bp were discarded.

| Treatment | Biological replicate | RIN value | # paired-end reads post trim (M) | % paired-end reads retained post trim | Avg. quality per read post trim | Q30 post trim |
|---|---|---|---|---|---|---|
| Ta_CTRL_0h | 1 | 6.2 | 9.92 | 75% | 35.6 | 98.6% |
| Ta_CTRL_0h | 2 | 7 | 15.96 | 99.2% | 35.47 | 97.7% |
| Ta_CTRL_0h | 3 | 8.8 | 10.22 | 99.4% | 35.53 | 98.3% |
| Ta_ECC_6h | 1 | 6.2 | 14.41 | 99.7% | 35.58 | 98.6% |
| Ta_ECC_6h | 2 | 7 | 11.13 | 99.2% | 35.45 | 97.6% |
| Ta_ECC_6h | 3 | 8.6 | 10.49 | 99% | 35.53 | 98.2% |
| Ta_SA1_6h | 1 | 6.7 | 10.62 | 99.5% | 35.55 | 98.5% |
| Ta_SA1_6h | 2 | 7.5 | 13.38 | 99.3% | 35.45 | 97.6% |
| Ta_SA1_6h | 3 | 8.6 | 10.41 | 98.5% | 35.45 | 97.7% |
| Ta_SAMv2_6h | 1 | 6.7 | 14.5 | 99.4% | 35.45 | 97.7% |
| Ta_SAMv2_6h | 2 | 8.9 | 10.15 | 98.7% | 35.53 | 98.2% |
| Ta_SAMv2_6h | 3 | 9 | 9.47 | 98.8% | 35.33 | 97.7% |
| Ta_SA5_6h | 1 | 6.3 | 12.49 | 99.5% | 35.53 | 98.4% |
| Ta_SA5_6h | 2 | 7 | 13.01 | 99.3% | 35.47 | 97.7% |
| Ta_SA5_6h | 3 | 8.9 | 10.15 | 98.7% | 35.37 | 97.6% |

**Table S5**. Significant differentially expressed *Diuraphis noxia* unigenes following a host shift from *Triticum aestivum* cultivar Gamtoos-S (0h) to near-isogenic, *Dn7* containing Gamtoos-R. Sampling occurred 6 h after aphid transfer (6h). Contrasts tested to determine effect of host shift in transcriptional regulation: DnSA1_6h - DnSA1_0h, DnSA5_6h - DnSA5_0h, DnSAMv2_6h - DnSAMv2_0h. NA, no blastp or InterProScan results. Sorted by FDR adjusted *p*-value.

| Target | Product description | Contrast | Adj. *p*-value | log2FC |
|---|---|---|---|---|
| GG_1409_c54_g6 | Solute carrier family 26 member 10-like | (DnSA1_6h + DnSAMv2_6h)/2-(DnSA1_0h + DnSAMv2_0h)/2 | 2.14E-04 | -1.024 |
| GG_711_c91_g1 | DNA-directed RNA polymerase II subunit RPB1-like | (DnSA1_6h + DnSAMv2_6h)/2-(DnSA1_0h + DnSAMv2_0h)/2 | 3.50E-03 | 1.054 |
| GG_1409_c54_g6 | Solute carrier family 26 member 10-like | DnSAMv2_6h - DnSAMv2_0h | 8.38E-03 | -1.137 |
| GG_585_c76_g2 | PREDICTED: uncharacterized protein LOC107172976 | (DnSA1_6h + DnSAMv2_6h)/2-(DnSA1_0h + DnSAMv2_0h)/2 | 1.28E-02 | 1.100 |
| GG_328_c18_g1 | DUF4806 domain-containing protein | (DnSA1_6h + DnSAMv2_6h)/2-(DnSA1_0h + DnSAMv2_0h)/2 | 2.30E-02 | -3.348 |
| GG_273_c58_g1 | Transposable element P transposase | (DnSA1_6h + DnSAMv2_6h)/2-(DnSA1_0h + DnSAMv2_0h)/2 | 2.41E-02 | 3.655 |
| GG_1562_c83_g1 | Facilitated trehalose transporter Tret1-like | DnSAMv2_6h - DnSAMv2_0h | 2.72E-02 | -1.035 |
| GG_1018_c0_g1 | DUF4806 domain-containing protein | (DnSA1_6h + DnSAMv2_6h)/2-(DnSA1_0h + DnSAMv2_0h)/2 | 2.97E-02 | 3.415 |
| GG_1534_c110_g1 | ---Na--- | (DnSA1_6h + DnSAMv2_6h)/2-(DnSA1_0h + DnSAMv2_0h)/2 | 3.09E-02 | -2.519 |
| GG_586_c11_g1 | Uncharacterized protein LOC111034199, partial | DnSAMv2_6h - DnSAMv2_0h | 3.16E-02 | 1.086 |
| GG_716_c92_g1 | Ankyrin repeat, PH and SEC7 domain containing protein secg-like | DnSA1_6h - DnSA1_0h | 3.22E-02 | -1.340 |
| GG_1409_c4_g1 | Venom protease-like | DnSA1_6h - DnSA1_0h | 3.39E-02 | -1.667 |
| GG_711_c151_g1 | Dynein axonemal heavy chain 5 | (DnSA1_6h + DnSAMv2_6h)/2-(DnSA1_0h + DnSAMv2_0h)/2 | 3.62E-02 | 1.169 |
| GG_585_c76_g2 | PREDICTED: uncharacterized protein LOC107172976 | DnSA1_6h - DnSA1_0h | 3.69E-02 | 1.703 |
| GG_586_c213_g1 | DDE 3 domain-containing protein | (DnSA1_6h + DnSAMv2_6h)/2-(DnSA1_0h + DnSAMv2_0h)/2 | 4.89E-02 | -4.156 |

**Table S6**. Significant differentially expressed *Diuraphis noxia* unigenes between avirulent biotype SA1 and its virulent direct progenitor, SAMv2. Aphids were placed on *Triticum aestivum* cultivar Gamtoos-S for one week (0h) whereafter it was transferred and contained on the near-isogenic line Gamtoos-R (containing the *D. noxia* resistance gene, *Dn7*; 6h). Contrasts tested: DnSAMv2_0h - DnSA1_0h, DnSAMv2_6h - DnSA1_6h, (DnSAMv2_0h + DnSAMv2_6h)/2 - (DnSA1_0h + DnSA1_6h)/2. Sorted by FDR adjusted *p*-value.

| **Target** | **Product description** | **Contrast** | **Adj. *p*-value** | **log2FC** |
|---|---|---|---|---|
| GG_1425_c31_g7 | ---NA--- | (DnSAMv2_0h + DnSAMv2_6h)/2-(DnSA1_0h + DnSA1_6h)/2 | 9.22E-03 | 1.034 |
| GG_1563_c7_g1 | Matrix metalloproteinase-16 | (DnSAMv2_0h + DnSAMv2_6h)/2-(DnSA1_0h + DnSA1_6h)/2 | 9.22E-03 | -1.068 |
| GG_1410_c151_g1 | Alpha-tocopherol transfer protein-like | (DnSAMv2_0h + DnSAMv2_6h)/2-(DnSA1_0h + DnSA1_6h)/2 | 1.56E-02 | -1.043 |
| GG_716_c74_g1 | Polycystic kidney disease and receptor for egg jelly-related protein | (DnSAMv2_0h + DnSAMv2_6h)/2-(DnSA1_0h + DnSA1_6h)/2 | 1.56E-02 | -1.005 |
| GG_1609_c1_g1 | Glucose dehydrogenase [FAD, quinone]-like | (DnSAMv2_0h + DnSAMv2_6h)/2-(DnSA1_0h + DnSA1_6h)/2 | 1.56E-02 | -1.082 |
| GG_713_c107_g3 | ---NA--- | (DnSAMv2_0h + DnSAMv2_6h)/2-(DnSA1_0h + DnSA1_6h)/2 | 1.56E-02 | 1.767 |
| GG_716_c40_g4 | Ribonuclease H-like domain, Ribonuclease H domain | (DnSAMv2_0h + DnSAMv2_6h)/2-(DnSA1_0h + DnSA1_6h)/2 | 1.82E-02 | -1.414 |
| GG_587_c1_g1 | Haemolymph juvenile hormone binding protein, JHBP | (DnSAMv2_0h + DnSAMv2_6h)/2-(DnSA1_0h + DnSA1_6h)/2 | 1.92E-02 | -1.078 |
| GG_1406_c61_g1 | Polyadenylate-binding protein 1-B-like | (DnSAMv2_0h + DnSAMv2_6h)/2-(DnSA1_0h + DnSA1_6h)/2 | 1.92E-02 | -1.149 |
| GG_403_c15_g1 | Arylsulfatase B-like | (DnSAMv2_0h + DnSAMv2_6h)/2-(DnSA1_0h + DnSA1_6h)/2 | 1.92E-02 | -1.012 |
| GG_587_c827_g1 | Cytoplasmic dynein 2 heavy chain 1 | (DnSAMv2_0h + DnSAMv2_6h)/2-(DnSA1_0h + DnSA1_6h)/2 | 1.92E-02 | 1.095 |
| GG_680_c325_g1 | ---NA--- | (DnSAMv2_0h + DnSAMv2_6h)/2-(DnSA1_0h + DnSA1_6h)/2 | 2.10E-02 | 2.531 |
| GG_757_c169_g1 | Uncharacterized protein LOC111030563 | (DnSAMv2_0h + DnSAMv2_6h)/2-(DnSA1_0h + DnSA1_6h)/2 | 2.10E-02 | 4.120 |

| | | | | |
|---|---|---|---|---|
| GG_1071_c16_g1 | Vasodilator-stimulated phosphoprotein-like | DnSAMv2_6h - DnSA1_6h | 2.70E-02 | -1.101 |
| GG_1071_c16_g2 | ---NA--- | DnSAMv2_6h - DnSA1_6h | 2.84E-02 | -1.153 |
| GG_1449_c28_g1 | Cysteine sulfinic acid decarboxylase | DnSAMv2_6h - DnSA1_6h | 2.84E-02 | -1.274 |
| GG_1563_c7_g1 | Matrix metalloproteinase-16 | DnSAMv2_6h - DnSA1_6h | 2.84E-02 | -1.224 |
| GG_1406_c335_g1 | PREDICTED: uncharacterized protein LOC107168679 | (DnSAMv2_0h + DnSAMv2_6h)/2-(DnSA1_0h + DnSA1_6h)/2 | 2.96E-02 | 1.348 |
| GG_1410_c101_g5 | ---NA--- | (DnSAMv2_0h + DnSAMv2_6h)/2-(DnSA1_0h + DnSA1_6h)/2 | 3.49E-02 | 1.633 |
| GG_216_c133_g1 | Calcium-binding mitochondrial carrier protein scamc-1-like | (DnSAMv2_0h + DnSAMv2_6h)/2-(DnSA1_0h + DnSA1_6h)/2 | 3.62E-02 | 2.861 |
| GG_273_c432_g1 | ---NA--- | (DnSAMv2_0h + DnSAMv2_6h)/2-(DnSA1_0h + DnSA1_6h)/2 | 3.92E-02 | 1.919 |
| GG_1609_c57_g1 | Nuclear control of atpase protein 2-like | (DnSAMv2_0h + DnSAMv2_6h)/2-(DnSA1_0h + DnSA1_6h)/2 | 4.17E-02 | 3.623 |
| GG_110_c4_g1 | Reverse transcriptase domain | DnSAMv2_6h - DnSA1_6h | 4.30E-02 | -1.015 |
| GG_1412_c12_g1 | Tyrosine 3-monooxygenase | DnSAMv2_6h - DnSA1_6h | 4.30E-02 | -1.359 |
| GG_1535_c2_g1 | Fatty acyl-CoA reductase wat-like | DnSAMv2_6h - DnSA1_6h | 4.30E-02 | -1.215 |
| GG_1625_c253_g1 | Retrovirus-related Pol polyprotein from type-1 retrotransposable element R1 2 | DnSAMv2_6h - DnSA1_6h | 4.30E-02 | -1.057 |
| GG_403_c15_g1 | Arylsulfatase B-like | DnSAMv2_6h - DnSA1_6h | 4.30E-02 | -1.420 |
| GG_587_c22_g2 | Protein spinster homolog 1 | DnSAMv2_6h - DnSA1_6h | 4.30E-02 | -1.266 |
| GG_598_c59_g6 | Uncharacterized protein LOC111030384 | DnSAMv2_6h - DnSA1_6h | 4.30E-02 | -1.005 |
| GG_711_c88_g1 | Fatty acyl-CoA reductase wat-like | DnSAMv2_6h - DnSA1_6h | 4.30E-02 | -1.104 |
| GG_587_c76_g3 | Progestin and adipoq receptor family member 3-like | DnSAMv2_6h - DnSA1_6h | 4.51E-02 | -1.023 |
| GG_634_c98_g1 | Cytochrome P450 4C1-like | DnSAMv2_6h - DnSA1_6h | 4.51E-02 | -2.509 |
| GG_716_c74_g1 | Polycystic kidney disease and receptor for egg jelly-related protein | DnSAMv2_6h - DnSA1_6h | 4.51E-02 | -1.065 |

| | | | | |
|---|---|---|---|---|
| GG_1044_c23_g4 | ---NA--- | (DnSAMv2_0h + DnSAMv2_6h)/2-(DnSA1_0h + DnSA1_6h)/2 | 4.52E-02 | 1.668 |
| GG_1535_c193_g1 | RNA-directed DNA polymerase from mobile element jockey-like | (DnSAMv2_0h + DnSAMv2_6h)/2-(DnSA1_0h + DnSA1_6h)/2 | 4.52E-02 | 1.544 |
| GG_1406_c61_g1 | Polyadenylate-binding protein 1-B-like | DnSAMv2_6h - DnSA1_6h | 4.64E-02 | -1.486 |
| GG_587_c1_g1 | ZZ-type zinc finger-containing protein 3 | DnSAMv2_6h - DnSA1_6h | 4.64E-02 | -1.343 |
| GG_1410_c151_g1 | Alpha-tocopherol transfer protein-like | DnSAMv2_6h - DnSA1_6h | 4.67E-02 | -1.161 |
| GG_1026_c26_g1 | L-xylulose reductase-like | (DnSAMv2_0h + DnSAMv2_6h)/2-(DnSA1_0h + DnSA1_6h)/2 | 4.81E-02 | 1.792 |
| GG_235_c1_g1 | PREDICTED: uncharacterized protein LOC107173559 | DnSAMv2_6h - DnSA1_6h | 4.81E-02 | -1.136 |

**Table S7**. Significant differentially expressed *Diuraphis noxia* unigenes between biotype SA5 and the closely related SA1 and SAMv2. Aphids were placed on *Triticum aestivum* cultivar Gamtoos-S for one week (0h) whereafter it was transferred and contained on the near-isogenic line Gamtoos-R (containing the *D. noxia* resistance gene, *Dn7*) for 6 h. Sorted by FDR adjusted *p*-value.

| **Target** | **Product description** | **Contrast** | **Adj. *p*-value** | **log2FC** |
|---|---|---|---|---|
| GG_1410_c331_g4 | Ring canal kelch protein | DnSA5_0h - DnSAMv2_0h | 7.56E-05 | 3.680 |
| GG_1410_c331_g4 | Ring canal kelch protein | DnSA5_0h - DnSA1_0h | 2.23E-04 | 4.331 |
| GG_1410_c331_g4 | Ring canal kelch protein | DnSA5_6h - DnSAMv2_6h | 2.44E-04 | 3.170 |
| GG_1581_c12_g1 | THAP domain-containing protein 1 B-like | DnSA5_0h - DnSA1_0h | 7.78E-04 | 1.352 |
| GG_1534_c108_g2 | Integrase catalytic domain-containing protein | DnSA5_0h - DnSA1_0h | 7.78E-04 | 2.119 |
| GG_1581_c12_g1 | THAP domain-containing protein 1 B-like | DnSA5_0h - DnSAMv2_0h | 1.01E-03 | 1.298 |
| GG_1575_c58_g1 | Retrovirus-related Pol polyprotein from transposon 17.6 | DnSA5_6h - DnSAMv2_6h | 1.18E-03 | -1.212 |
| GG_1410_c331_g4 | Ring canal kelch protein | DnSA5_6h - DnSA1_6h | 1.31E-03 | 2.848 |
| GG_1575_c58_g1 | Retrovirus-related Pol polyprotein from transposon 17.6 | DnSA5_6h - DnSA1_6h | 2.17E-03 | -1.180 |
| GG_7_c1_g1 | ---NA--- | DnSA5_6h - DnSA1_6h | 3.75E-03 | -1.439 |
| GG_1575_c58_g1 | Retrovirus-related Pol polyprotein from transposon 17.6 | DnSA5_0h - DnSA1_0h | 3.92E-03 | -1.135 |
| GG_1534_c108_g2 | Integrase catalytic domain-containing protein | DnSA5_0h - DnSAMv2_0h | 4.50E-03 | 1.606 |
| GG_585_c198_g1 | Histone acetyltransferase KAT6B | DnSA5_0h - DnSAMv2_0h | 4.50E-03 | 5.138 |
| GG_1581_c12_g1 | THAP domain-containing protein 1 B-like | DnSA5_6h - DnSAMv2_6h | 5.28E-03 | 1.043 |
| GG_1534_c108_g2 | Integrase catalytic domain-containing protein | DnSA5_6h - DnSAMv2_6h | 5.28E-03 | 1.609 |
| GG_1575_c58_g1 | Retrovirus-related Pol polyprotein from transposon 17.6 | DnSA5_0h - DnSAMv2_0h | 5.29E-03 | -1.058 |
| GG_713_c29_g1 | Protein ALP1-like | DnSA5_0h - DnSA1_0h | 6.72E-03 | 1.047 |
| GG_1535_c33_g4 | AP2/ERF domain-containing protein PFD0985w-like | DnSA5_6h - DnSAMv2_6h | 1.05E-02 | -1.276 |
| GG_585_c198_g1 | Histone acetyltransferase KAT6B | DnSA5_0h - DnSA1_0h | 1.10E-02 | 5.122 |
| GG_676_c101_g1 | Hypothetical protein AGLY_013878 | DnSA5_6h - DnSAMv2_6h | 1.15E-02 | -4.844 |

| | | | | |
|---|---|---|---|---|
| GG_676_c101_g1 | Hypothetical protein AGLY_013878 | DnSA5_6h - DnSA1_6h | 1.16E-02 | -4.790 |
| GG_716_c4_g3 | Integrase catalytic domain-containing protein | DnSA5_0h - DnSA1_0h | 1.55E-02 | 1.317 |
| GG_7_c1_g1 | ---NA--- | DnSA5_0h - DnSAMv2_0h | 1.58E-02 | -1.395 |
| GG_1564_c179_g1 | ---NA--- | DnSA5_6h - DnSA1_6h | 1.82E-02 | -1.003 |
| GG_7_c1_g1 | ---NA--- | DnSA5_6h - DnSAMv2_6h | 1.83E-02 | -1.216 |
| GG_711_c195_g1 | Uncharacterized protein FWK35_00022957 | DnSA5_6h - DnSAMv2_6h | 2.03E-02 | -1.491 |
| GG_585_c198_g1 | Histone acetyltransferase KAT6B | DnSA5_6h - DnSAMv2_6h | 2.48E-02 | 3.869 |
| GG_1534_c108_g2 | Integrase catalytic domain-containing protein | DnSA5_6h - DnSA1_6h | 2.61E-02 | 1.282 |
| GG_711_c195_g1 | Uncharacterized protein FWK35_00022957 | DnSA5_6h - DnSA1_6h | 2.95E-02 | -1.412 |
| GG_130_c65_g1 | DDE 3 domain-containing protein | DnSA5_6h - DnSAMv2_6h | 3.61E-02 | -3.525 |
| GG_585_c353_g1 | ---NA--- | DnSA5_0h - DnSA1_0h | 4.43E-02 | 3.989 |
| GG_130_c65_g1 | DDE 3 domain-containing protein | DnSA5_0h - DnSA1_0h | 4.48E-02 | -5.665 |
| GG_7_c1_g1 | ---NA--- | DnSA5_0h - DnSA1_0h | 4.48E-02 | -1.263 |
| GG_1425_c99_g2 | MULE domain-containing protein | DnSA5_0h - DnSA1_0h | 4.58E-02 | 1.737 |
| GG_130_c65_g1 | DDE 3 domain-containing protein | DnSA5_0h - DnSAMv2_0h | 4.89E-02 | -5.686 |
| GG_1425_c31_g7 | ---NA--- | DnSA5_0h - DnSAMv2_0h | 4.89E-02 | -1.230 |
| GG_585_c353_g1 | ---NA--- | DnSA5_0h - DnSAMv2_0h | 4.89E-02 | 3.417 |

**Table S8**. Significant differential alternative splicing (DAS) of *Diuraphis noxi*a unigenes in all contrasts tested. Aphids were placed on *Triticum aestivum* cultivar Gamtoos-S for one week (0h) whereafter it was transferred and contained on the near-isogenic line Gamtoos-R (containing the *D. noxia* resistance gene, *Dn7*) for 6 h (6h). The Limma voomWithQualityWeights function (Liu *et al.*, 2015; Ritchie *et al.*, 2015) was used to calculated DAS by comparing the change of each transcript to the gene change in gene expression. The *p*-values are then combined to a single gene-level *p*-value using an F-test. Top 50 sorted by adjusted *p*-value included here. PS, percent spliced.

| **Target** | **Product description** | **Contrast** | **Adj. *p*-value** | **Maxdelt aPS** | **log2 FC** |
|---|---|---|---|---|---|
| GG_1410_c233_g1 | Glycoprotein-N-acetylgalactosamine 3-beta-galactosyltransferase 1-like | DnSA5_6h - DnSAMv2_6h | 8.00E-12 | 0.112 | -0.100 |
| GG_1575_c12_g1 | 52 kDa repressor of the inhibitor of the protein kinase-like | DnSA5_6h - DnSA1_6h | 2.68E-10 | -0.161 | -0.047 |
| GG_1575_c12_g1 | 52 kDa repressor of the inhibitor of the protein kinase-like | DnSA5_6h - DnSAMv2_6h | 2.47E-06 | -0.106 | -0.099 |
| GG_1575_c12_g1 | 52 kDa repressor of the inhibitor of the protein kinase-like | DnSA5_0h - DnSAMv2_0h | 6.16E-06 | -0.118 | 0.011 |
| GG_1581_c7_g1 | MAGUK p55 subfamily member 7 | DnSA5_6h - DnSA1_6h | 2.61E-05 | -0.281 | -0.024 |
| GG_716_c46_g1 | Lactoperoxidase | DnSA5_6h - DnSAMv2_6h | 5.21E-05 | -0.269 | 0.117 |
| GG_716_c46_g1 | Lactoperoxidase | DnSA5_0h - DnSAMv2_0h | 7.07E-05 | 0.384 | 0.097 |
| GG_1581_c7_g1 | MAGUK p55 subfamily member 7 | (DnSAMv2_0h + DnSAMv2_6h)/2-(DnSA1_0h + DnSA1_6h)/2 | 1.88E-04 | 0.164 | 0.050 |
| GG_716_c54_g1 | ---NA--- | DnSA5_6h - DnSA1_6h | 4.25E-04 | 0.936 | -0.256 |
| GG_1409_c78_g1 | Activator of 90 kDa heat shock protein ATPase homolog 1 | DnSA5_0h - DnSAMv2_0h | 7.04E-04 | -0.225 | 0.088 |
| GG_1575_c26_g2 | RNA-directed DNA polymerase from mobile element jockey | DnSA5_6h - DnSAMv2_6h | 7.07E-04 | -0.855 | -1.536 |
| GG_716_c54_g1 | ---NA--- | DnSA5_6h - DnSAMv2_6h | 7.07E-04 | 0.892 | -0.165 |
| GG_1425_c54_g1 | Toll-interacting protein-like | DnSAMv2_6h - DnSA1_6h | 9.47E-04 | 0.231 | -0.061 |
| GG_1575_c93_g1 | Inositol polyphosphate 1-phosphatase | DnSA5_6h - DnSA1_6h | 9.53E-04 | -0.545 | 0.308 |
| GG_716_c46_g1 | Lactoperoxidase | DnSA5_6h - DnSA1_6h | 9.53E-04 | -0.252 | -0.087 |
| GG_1575_c93_g1 | Inositol polyphosphate 1-phosphatase | DnSA5_6h - DnSAMv2_6h | 9.85E-04 | -0.509 | 0.373 |
| GG_716_c54_g1 | ---NA--- | DnSA5_0h - DnSAMv2_0h | 1.01E-03 | 0.662 | -0.054 |
| GG_716_c46_g1 | Lactoperoxidase | DnSA5_0h - DnSA1_0h | 1.09E-03 | 0.384 | 0.171 |

| | | | | | |
|---|---|---|---|---|---|
| GG_1575_c18_g1 | Zinc finger protein 484-like | DnSA5_6h - DnSA1_6h | 1.16E-03 | 0.266 | 0.007 |
| GG_728_c3_g1 | Protein ALP1-like | DnSA5_6h - DnSAMv2_6h | 1.19E-03 | 0.473 | 0.607 |
| GG_721_c102_g1 | ADP-ribosylation factor 1 | DnSAMv2_0h - DnSA1_0h | 1.38E-03 | 0.218 | 0.072 |
| GG_623_c27_g1 | Cyclin-dependent kinase 1-like | DnSA5_6h - DnSA1_6h | 1.64E-03 | -0.240 | 0.125 |
| GG_728_c3_g1 | Protein ALP1-like | DnSA5_6h - DnSA1_6h | 1.64E-03 | -0.365 | 0.172 |
| GG_1052_c9_g1 | Tubulin polyglutamylase TTLL5 | DnSA5_6h - DnSA5_0h | 1.67E-03 | 0.212 | 0.164 |
| GG_603_c4_g1 | Protein O-GlcNAcase | DnSA5_6h - DnSAMv2_6h | 1.77E-03 | 0.296 | -0.037 |
| GG_1575_c93_g1 | Inositol polyphosphate 1-phosphatase | DnSA5_0h - DnSA1_0h | 1.77E-03 | -0.697 | 0.662 |
| GG_266_c17_g1 | ---NA--- | DnSAMv2_6h - DnSA1_6h | 2.15E-03 | 0.549 | -0.339 |
| GG_1449_c22_g1 | Protein sel-1 homolog 1 | DnSA5_6h - DnSA5_0h | 2.23E-03 | 0.138 | -0.049 |
| GG_737_c62_g1 | Guanine nucleotide exchange factor subunit Rich | DnSA5_6h - DnSAMv2_6h | 2.44E-03 | -0.108 | -0.036 |
| GG_587_c115_g1 | Ankyrin repeat and sterile alpha motif domain-containing protein 1B-like | DnSAMv2_0h - DnSA1_0h | 2.68E-03 | -0.121 | 0.035 |
| GG_603_c4_g1 | Protein O-GlcNAcase | DnSA5_0h - DnSAMv2_0h | 2.79E-03 | 0.339 | -0.050 |
| GG_1415_c78_g1 | PREDICTED: uncharacterized protein LOC107169683 | DnSA5_6h - DnSA1_6h | 2.80E-03 | 0.219 | 0.047 |
| GG_720_c4_g1 | Centrosomal and chromosomal factor | DnSA5_6h - DnSA1_6h | 2.80E-03 | -0.380 | -0.017 |
| GG_1448_c17_g1 | Monocarboxylate transporter 12-like | DnSA5_0h - DnSAMv2_0h | 2.96E-03 | -0.191 | 0.277 |
| GG_1562_c80_g1 | Protein apterous-like | DnSA5_0h - DnSAMv2_0h | 2.96E-03 | 0.454 | -0.116 |
| GG_347_c4_g2 | ---NA--- | (DnSA1_6h + DnSAMv2_6h)/2-(DnSA1_0h + DnSAMv2_0h)/2 | 2.96E-03 | -0.166 | -0.123 |
| GG_598_c27_g1 | Supervillin | DnSA5_6h - DnSAMv2_6h | 3.12E-03 | 0.104 | -0.174 |
| GG_720_c4_g1 | Centrosomal and chromosomal factor | DnSA5_6h - DnSAMv2_6h | 3.12E-03 | -0.347 | -0.026 |
| GG_672_c7_g1 | Uncharacterized protein FWK35_00018137 | DnSA5_0h - DnSAMv2_0h | 3.24E-03 | -0.430 | -0.159 |
| GG_716_c83_g1 | Mannose-6-phosphate isomerase | DnSA5_6h - DnSAMv2_6h | 3.40E-03 | 0.205 | -0.195 |
| GG_1071_c24_g3 | NADH dehydrogenase (ubiquinone) complex I, assembly factor 6 | (DnSAMv2_0h + DnSAMv2_6h)/2-(DnSA1_0h + DnSA1_6h)/2 | 3.48E-03 | -0.312 | -0.042 |
| GG_859_c93_g1 | Trimethylguanosine synthase-like | DnSA5_0h - DnSAMv2_0h | 3.67E-03 | -0.143 | -0.063 |
| GG_713_c119_g1 | Membrane-bound alkaline phosphatase-like | DnSA5_6h - DnSA1_6h | 3.95E-03 | -0.146 | -0.164 |

| | | | | | |
|---|---|---|---|---|---|
| GG_1575_c30_g1 | Sushi, von Willebrand factor type A, EGF and pentraxin domain-containing protein 1 | DnSA5_6h - DnSAMv2_6h | 4.19E-03 | 0.232 | 0.048 |
| GG_1409_c9_g1 | Golgi integral membrane protein 4-like | DnSAMv2_6h - DnSAMv2_0h | 4.33E-03 | -0.106 | 0.034 |
| GG_716_c21_g1 | GILT-like protein 1 | DnSA5_6h - DnSA1_6h | 4.43E-03 | 0.220 | -0.019 |
| GG_1425_c54_g1 | Toll-interacting protein-like | DnSA5_6h - DnSA1_6h | 5.02E-03 | 0.115 | 0.018 |
| GG_1575_c445_g1 | Probable lysine-specific demethylase 4B | DnSA5_0h - DnSAMv2_0h | 5.52E-03 | 0.505 | 0.032 |
| GG_1575_c93_g1 | Inositol polyphosphate 1-phosphatase | DnSA5_0h - DnSAMv2_0h | 5.52E-03 | -0.556 | 0.695 |
| GG_672_c3_g1 | Zinc finger CCCH domain-containing protein 11A-like | DnSA5_0h - DnSAMv2_0h | 5.52E-03 | -0.249 | -0.101 |

**Table S9**. Significant differentially expressed genes in *Triticum aestivum* cultivar Gamtoos-R (containing the *Dn7*) between the 0 h control and samples taken at 6 h of aphid containment or empty clip cage (ECC). Contrasts tested: CTRL 6 h - Ta_CTRL_0h, SA1 6 h - Ta_CTRL_0h, SAMv2 6 h - Ta_CTRL_0h and SA5 6 h - Ta_CTRL_0h. CTRL 6 h. Top 50 sorted by FDR adjusted *p*-value.

| **Target** | **Product description** | **Contrast** | **Adj. *p*-value** | **log2FC** |
|---|---|---|---|---|
| LOC123127760 | Two-component response regulator-like PRR1 | Ta_SAMv2_6h - Ta_CTRL_0h | 7.22E-06 | 4.036 |
| LOC123137489 | Two-component response regulator-like PRR1 | Ta_SAMv2_6h - Ta_CTRL_0h | 7.22E-06 | 3.783 |
| LOC123162779 | Zinc finger protein CONSTANS-LIKE 9-like | Ta_SAMv2_6h - Ta_CTRL_0h | 7.22E-06 | 2.939 |
| LOC123190226 | Protein LNK4-like | Ta_SA5_6h - Ta_CTRL_0h | 8.31E-06 | -4.876 |
| LOC123127760 | Two-component response regulator-like PRR1 | Ta_SA5_6h - Ta_CTRL_0h | 8.31E-06 | 3.985 |
| LOC123137489 | Two-component response regulator-like PRR1 | Ta_SA5_6h - Ta_CTRL_0h | 8.31E-06 | 3.653 |
| LOC123162779 | Zinc finger protein CONSTANS-LIKE 9-like | Ta_SA5_6h - Ta_CTRL_0h | 8.31E-06 | 3.085 |
| LOC123162779 | Zinc finger protein CONSTANS-LIKE 9-like | Ta_SA1_6h - Ta_CTRL_0h | 8.42E-06 | 3.137 |
| LOC123086549 | NAD(P)H-quinone oxidoreductase subunit T, chloroplastic-like | Ta_SAMv2_6h - Ta_CTRL_0h | 1.33E-05 | -3.658 |
| LOC123127760 | Two-component response regulator-like PRR1 | Ta_ECC_6h - Ta_CTRL_0h | 1.35E-05 | 3.892 |
| LOC123137489 | Two-component response regulator-like PRR1 | Ta_ECC_6h - Ta_CTRL_0h | 1.35E-05 | 3.651 |
| LOC123162779 | Zinc finger protein CONSTANS-LIKE 9-like | Ta_ECC_6h - Ta_CTRL_0h | 1.35E-05 | 2.740 |
| LOC123167272 | Zinc finger protein CONSTANS-LIKE 9-like | Ta_SA5_6h - Ta_CTRL_0h | 1.47E-05 | 2.776 |
| LOC123167076 | Protein LHY-like | Ta_ECC_6h - Ta_CTRL_0h | 2.16E-05 | -5.271 |
| LOC123190226 | Protein LNK4-like | Ta_ECC_6h - Ta_CTRL_0h | 2.16E-05 | -4.125 |
| LOC123167272 | Zinc finger protein CONSTANS-LIKE 9-like | Ta_SAMv2_6h - Ta_CTRL_0h | 2.39E-05 | 2.658 |
| LOC123190226 | Protein LNK4-like | Ta_SAMv2_6h - Ta_CTRL_0h | 2.86E-05 | -4.041 |
| LOC123167272 | Zinc finger protein CONSTANS-LIKE 9-like | Ta_SA1_6h - Ta_CTRL_0h | 2.87E-05 | 2.823 |
| LOC123137489 | Two-component response regulator-like PRR1 | Ta_SA1_6h - Ta_CTRL_0h | 3.14E-05 | 3.364 |
| LOC123127760 | Two-component response regulator-like PRR1 | Ta_SA1_6h - Ta_CTRL_0h | 3.30E-05 | 3.454 |

| | | | | |
|---|---|---|---|---|
| LOC123167272 | Zinc finger protein CONSTANS-LIKE 9-like | Ta_ECC_6h - Ta_CTRL_0h | 4.48E-05 | 2.471 |
| LOC123190226 | Protein LNK4-like | Ta_SA1_6h - Ta_CTRL_0h | 4.52E-05 | -3.953 |
| LOC123167076 | Protein LHY-like | Ta_SAMv2_6h - Ta_CTRL_0h | 4.56E-05 | -4.853 |
| LOC100682474 | Protein LHY | Ta_ECC_6h - Ta_CTRL_0h | 4.80E-05 | -5.255 |
| LOC123086549 | NAD(P)H-quinone oxidoreductase subunit T, chloroplastic-like | Ta_ECC_6h - Ta_CTRL_0h | 4.80E-05 | -3.074 |
| LOC123045606 | Probable protein phosphatase 2C 40 | Ta_SA5_6h - Ta_CTRL_0h | 5.25E-05 | 3.295 |
| LOC123045606 | Probable protein phosphatase 2C 40 | Ta_SAMv2_6h - Ta_CTRL_0h | 5.45E-05 | 3.242 |
| LOC123187442 | Cyclic dof factor 1-like | Ta_ECC_6h - Ta_CTRL_0h | 5.58E-05 | -3.586 |
| LOC100682474 | Protein LHY | Ta_SAMv2_6h - Ta_CTRL_0h | 5.88E-05 | -5.177 |
| LOC123114430 | Beta-fructofuranosidase, insoluble isoenzyme 4-like | Ta_SA1_6h - Ta_CTRL_0h | 6.59E-05 | 4.247 |
| LOC123150352 | Zinc finger protein CONSTANS-LIKE 9-like | Ta_SA1_6h - Ta_CTRL_0h | 6.59E-05 | 3.231 |
| LOC123045606 | Probable protein phosphatase 2C 40 | Ta_SA1_6h - Ta_CTRL_0h | 6.59E-05 | 3.168 |
| LOC123105964 | Probable aquaporin TIP4-1 | Ta_SA1_6h - Ta_CTRL_0h | 6.59E-05 | 2.978 |
| LOC123114430 | Beta-fructofuranosidase, insoluble isoenzyme 4-like | Ta_ECC_6h - Ta_CTRL_0h | 7.11E-05 | 4.199 |
| LOC123191968 | Protein SYM1-like | Ta_ECC_6h - Ta_CTRL_0h | 7.11E-05 | -2.110 |
| LOC123167076 | Protein LHY-like | Ta_SA5_6h - Ta_CTRL_0h | 7.27E-05 | -4.546 |
| LOC123114430 | Beta-fructofuranosidase, insoluble isoenzyme 4-like | Ta_SA5_6h - Ta_CTRL_0h | 7.27E-05 | 4.226 |
| LOC123091549 | Root phototropism protein 2-like | Ta_SA5_6h - Ta_CTRL_0h | 7.27E-05 | -4.092 |
| LOC123062854 | Uncharacterized protein LOC123062854 | Ta_SA5_6h - Ta_CTRL_0h | 7.27E-05 | -3.424 |
| LOC123150352 | Zinc finger protein CONSTANS-LIKE 9-like | Ta_SA5_6h - Ta_CTRL_0h | 7.27E-05 | 3.135 |
| LOC123105964 | Probable aquaporin TIP4-1 | Ta_SA5_6h - Ta_CTRL_0h | 7.27E-05 | 2.981 |
| LOC123086549 | NAD(P)H-quinone oxidoreductase subunit T, chloroplastic-like | Ta_SA5_6h - Ta_CTRL_0h | 7.27E-05 | -2.917 |
| LOC123143620 | Probable protein phosphatase 2C 71 | Ta_SA5_6h - Ta_CTRL_0h | 7.27E-05 | -1.860 |

| | | | | |
|---|---|---|---|---|
| LOC123114430 | Beta-fructofuranosidase, insoluble isoenzyme 4-like | Ta_SAMv2_6h - Ta_CTRL_0h | 7.81E-05 | 4.153 |
| LOC123143620 | Probable protein phosphatase 2C 71 | Ta_SAMv2_6h - Ta_CTRL_0h | 7.81E-05 | -1.833 |
| LOC123045606 | Probable protein phosphatase 2C 40 | Ta_ECC_6h - Ta_CTRL_0h | 8.12E-05 | 3.109 |
| LOC123150352 | Zinc finger protein CONSTANS-LIKE 9-like | Ta_SAMv2_6h - Ta_CTRL_0h | 8.27E-05 | 3.083 |
| LOC123094036 | Cold-regulated protein 27-like | Ta_SA5_6h - Ta_CTRL_0h | 8.46E-05 | 6.416 |
| LOC123086549 | NAD(P)H-quinone oxidoreductase subunit T, chloroplastic-like | Ta_SA1_6h - Ta_CTRL_0h | 8.56E-05 | -2.924 |
| LOC123105964 | Probable aquaporin TIP4-1 | Ta_SAMv2_6h - Ta_CTRL_0h | 8.81E-05 | 2.805 |

**Table S10**. Significant differentially expressed genes in *Triticum aestivum* cultivar Gamtoos-R (containing the *Dn7 Diuraphis noxia* resistance gene) between the 6 h control (empty clip cage; ECC) and plants on which aphid biotypes SA1, SAMv2 and SA5 were contained using clip cages. Sorted by FDR adjusted *p*-value.

| **Target** | **Product description** | **Contrast** | **Adj. *p*-value** | **log2FC** |
|---|---|---|---|---|
| LOC123121546 | Protein TIFY 10c-like | Ta_SA5_6h - Ta_ECC_6h | 9.23E-03 | 5.489 |
| LOC123175967 | Probable 1-deoxy-D-xylulose-5-phosphate synthase2, chloroplastic | Ta_SA5_6h - Ta_ECC_6h | 9.23E-03 | 3.328 |
| LOC123098267 | Probable 1-deoxy-D-xylulose-5-phosphate synthase2, chloroplastic | Ta_SA5_6h - Ta_ECC_6h | 9.23E-03 | 3.076 |
| LOC123181002 | Probable 1-deoxy-D-xylulose-5-phosphate synthase2, chloroplastic | Ta_SA5_6h - Ta_ECC_6h | 1.25E-02 | 3.089 |
| LOC123103761 | Protein TIFY 10c-like | Ta_SA5_6h - Ta_ECC_6h | 1.44E-02 | 5.036 |
| LOC123088283 | Bisdemethoxycurcumin synthase-like | Ta_SA5_6h - Ta_ECC_6h | 1.44E-02 | 4.789 |
| LOC123056948 | Mitogen-activated protein kinase kinase kinase 18-like | Ta_SA5_6h - Ta_ECC_6h | 1.44E-02 | 4.699 |
| LOC100873096 | Pathogenesis-related protein 1 | Ta_SA5_6h - Ta_ECC_6h | 1.44E-02 | 4.688 |
| LOC123187136 | Putative methyltransferase DDB_G0268948 | Ta_SA5_6h - Ta_ECC_6h | 1.44E-02 | 3.471 |
| LOC123046106 | Phenylalanine ammonia-lyase-like | Ta_SA5_6h - Ta_ECC_6h | 1.44E-02 | 2.560 |
| LOC123052098 | Protein TIFY 10b-like | Ta_SA5_6h - Ta_ECC_6h | 1.44E-02 | 1.529 |
| LOC123081537 | Protein TIFY 11b-like | Ta_SA5_6h - Ta_ECC_6h | 1.54E-02 | 4.727 |
| LOC123092461 | Jasmonate-induced oxygenase 1-like | Ta_SA5_6h - Ta_ECC_6h | 1.84E-02 | 4.151 |
| LOC123098267 | Probable 1-deoxy-D-xylulose-5-phosphate synthase2, chloroplastic | Ta_SA1_6h - Ta_ECC_6h | 2.42E-02 | 2.844 |
| LOC123188944 | Transcription factor BHLH6-like | Ta_SA5_6h - Ta_ECC_6h | 2.73E-02 | 4.185 |
| LOC123051893 | 4-coumarate--CoA ligase 5-like | Ta_SA5_6h - Ta_ECC_6h | 2.73E-02 | 2.790 |
| LOC123132353 | Putative linoleate 9S-lipoxygenase 3 | Ta_SA5_6h - Ta_ECC_6h | 2.97E-02 | 15.195 |
| LOC123130196 | S-(+)-linalool synthase, chloroplastic-like | Ta_SA5_6h - Ta_ECC_6h | 3.37E-02 | 5.353 |
| LOC123124828 | Zinc finger protein 2-like | Ta_SA5_6h - Ta_ECC_6h | 3.37E-02 | 4.667 |
| LOC123085113 | Allene oxide synthase 2-like | Ta_SA5_6h - Ta_ECC_6h | 3.37E-02 | 4.237 |

| | | | | |
|---|---|---|---|---|
| LOC123045198 | Transcription factor BHLH6-like | Ta_SA5_6h - Ta_ECC_6h | 3.37E-02 | 4.099 |
| LOC123098416 | Cytochrome b561 and DOMON domain-containing protein At5g47530-like | Ta_SA5_6h - Ta_ECC_6h | 3.37E-02 | 3.643 |
| LOC123083133 | Cytochrome P450 711A1-like | Ta_SA5_6h - Ta_ECC_6h | 3.37E-02 | 3.096 |
| LOC123167005 | Chaperone protein dnaJ 8, chloroplastic-like | Ta_SA5_6h - Ta_ECC_6h | 3.37E-02 | 2.994 |
| LOC123093777 | Pore-forming toxin-like protein Hfr-2 | Ta_SA5_6h - Ta_ECC_6h | 3.45E-02 | 6.708 |
| LOC123116183 | Protein TIFY 10c-like | Ta_SA5_6h - Ta_ECC_6h | 3.45E-02 | 4.623 |
| LOC123113260 | Transcription factor BHLH148-like | Ta_SA5_6h - Ta_ECC_6h | 3.77E-02 | 2.425 |
| LOC101290672 | CBL-interacting protein kinase 29 | Ta_SA5_6h - Ta_ECC_6h | 3.81E-02 | 2.044 |
| LOC123053034 | Transcription factor BHLH6-like | Ta_SA5_6h - Ta_ECC_6h | 4.01E-02 | 4.864 |
| LOC123138844 | S-(+)-linalool synthase, chloroplastic-like | Ta_SA5_6h - Ta_ECC_6h | 4.09E-02 | 5.293 |
| LOC123052522 | Probable glutathione S-transferase GSTU6 | Ta_SA5_6h - Ta_ECC_6h | 4.09E-02 | 4.009 |
| LOC123135793 | Aldo-keto reductase family 4 member C10-like | Ta_SA5_6h - Ta_ECC_6h | 4.56E-02 | 4.481 |
| LOC123121546 | Protein TIFY 10c-like | Ta_SA1_6h - Ta_ECC_6h | 4.61E-02 | 4.757 |
| LOC123056948 | Mitogen-activated protein kinase kinase kinase 18-like | Ta_SA1_6h - Ta_ECC_6h | 4.61E-02 | 4.720 |
| LOC123081537 | Protein TIFY 11b-like | Ta_SA1_6h - Ta_ECC_6h | 4.61E-02 | 4.587 |
| LOC123103761 | Protein TIFY 10c-like | Ta_SA1_6h - Ta_ECC_6h | 4.61E-02 | 4.584 |
| LOC100873096 | Pathogenesis-related protein 1 | Ta_SA1_6h - Ta_ECC_6h | 4.61E-02 | 4.303 |
| LOC123188944 | Transcription factor BHLH6-like | Ta_SA1_6h - Ta_ECC_6h | 4.61E-02 | 4.292 |
| LOC123181002 | Probable 1-deoxy-D-xylulose-5-phosphate synthase2, chloroplastic | Ta_SA1_6h - Ta_ECC_6h | 4.61E-02 | 2.632 |
| LOC123135793 | Aldo-keto reductase family 4 member C10-like | Ta_SA1_6h - Ta_ECC_6h | 4.68E-02 | 5.061 |
| LOC123045198 | Transcription factor BHLH6-like | Ta_SA1_6h - Ta_ECC_6h | 4.68E-02 | 4.284 |
| LOC123140864 | S-(+)-linalool synthase, chloroplastic-like | Ta_SA5_6h - Ta_ECC_6h | 4.70E-02 | 4.323 |

| | | | | |
|---|---|---|---|---|
| LOC123121546 | Protein TIFY 10c-like | Ta_SAMv2_6h - Ta_ECC_6h | 4.81E-02 | 5.170 |

**Table S11**. Significant differential alternative splicing (DAS) of genes in *Triticum aestivum* cultivar Gamtoos-R (containing *Dn7*) in all contrasts tested using the Limma voomWithQualityWeights function (Liu *et al.*, 2015; Ritchie *et al.*, 2015). DAS is calculated by comparing the change of each transcript to the gene change in gene expression. The *p*-values are then combined to a single gene-level *p*-value using an F-test. Top 50 sorted by FDR adjusted *p*-value. PS, percent spliced.

| Target | Product description | Contrast | Adj. *p*-val | Max delta PS | log2FC |
|---|---|---|---|---|---|
| LOC123057677 | Uncharacterized lipoprotein syc1174_c-like | Ta_SA5_6h - Ta_CTRL_0h | 3.26E-04 | -1.000 | -0.334 |
| LOC123057677 | Uncharacterized lipoprotein syc1174_c-like | Ta_SA5_6h - Ta_ECC_6h | 3.76E-04 | -1.000 | 0.230 |
| LOC123057677 | Uncharacterized lipoprotein syc1174_c-like | Ta_SAMv2_6h - Ta_SA5_6h | 3.85E-04 | 1.000 | -0.439 |
| LOC542897 | E3 ubiquitin-protein ligase RHF2A | Ta_SAMv2_6h - Ta_ECC_6h | 6.75E-04 | -0.340 | -0.047 |
| LOC123079583 | Probable serine/threonine-protein kinase PBL1 | Ta_SA5_6h - Ta_ECC_6h | 1.11E-03 | 0.975 | 1.025 |
| LOC123062011 | Probable complex I intermediate-associated protein 30 | Ta_SA5_6h - Ta_ECC_6h | 1.11E-03 | 0.558 | -0.290 |
| LOC123146363 | Protein argonaute 1C-like | Ta_ECC_6h - Ta_CTRL_0h | 1.16E-03 | 0.434 | -0.356 |
| LOC123091823 | Uncharacterized LOC123091823 | Ta_SAMv2_6h - Ta_SA1_6h | 1.33E-03 | -0.561 | -0.048 |
| LOC123103571 | ASC1-like protein 1 | Ta_SA1_6h - Ta_CTRL_0h | 1.33E-03 | 0.812 | 0.155 |
| LOC101669848 | Uncharacterized protein LOC101669848 | Ta_SA5_6h - Ta_CTRL_0h | 1.35E-03 | -0.684 | 0.137 |
| LOC123057677 | Uncharacterized lipoprotein syc1174_c-like | Ta_SA1_6h - Ta_ECC_6h | 1.35E-03 | -0.879 | 0.741 |
| LOC123062011 | Probable complex I intermediate-associated protein 30 | Ta_SA1_6h - Ta_ECC_6h | 1.35E-03 | 0.558 | -0.339 |
| LOC123070240 | Uncharacterized LOC123070240 | Ta_SA1_6h - Ta_ECC_6h | 1.35E-03 | -0.728 | -0.050 |
| LOC123057677 | Uncharacterized lipoprotein syc1174_c-like | Ta_SA1_6h - Ta_CTRL_0h | 1.56E-03 | -0.879 | 0.177 |
| LOC123103571 | ASC1-like protein 1 | Ta_SA1_6h - Ta_SA5_6h | 1.78E-03 | 0.775 | -0.048 |
| LOC123045258 | Phospholipid-transporting ATPase 2-like | Ta_SA5_6h - Ta_ECC_6h | 1.80E-03 | 1.000 | 0.184 |
| LOC123052739 | Uncharacterized protein LOC123052739 | Ta_SA5_6h - Ta_ECC_6h | 1.80E-03 | -0.422 | -0.109 |
| LOC123045258 | Phospholipid-transporting ATPase 2-like | Ta_SAMv2_6h - Ta_ECC_6h | 2.02E-03 | 1.000 | 0.560 |
| LOC123079583 | Probable serine/threonine-protein kinase PBL1 | Ta_ECC_6h - Ta_CTRL_0h | 2.09E-03 | -1.000 | -0.637 |
| LOC123057677 | Uncharacterized lipoprotein syc1174_c-like | Ta_SAMv2_6h - Ta_SA1_6h | 2.22E-03 | 0.879 | -0.950 |
| LOC123062011 | Probable complex I intermediate-associated protein 30 | Ta_SAMv2_6h - Ta_ECC_6h | 2.36E-03 | 0.558 | -0.634 |
| LOC123045075 | Uncharacterized protein LOC123045075 | Ta_SAMv2_6h - Ta_ECC_6h | 2.36E-03 | 0.676 | -0.456 |
| LOC123119727 | Boron transporter 1-like | Ta_SAMv2_6h - Ta_ECC_6h | 2.36E-03 | 0.450 | -0.352 |

| | | | | | |
|---|---|---|---|---|---|
| LOC123070901 | Uncharacterized protein LOC123070901 | Ta_SA5_6h - Ta_ECC_6h | 2.48E-03 | -0.221 | -0.055 |
| LOC123088266 | Uncharacterized protein LOC123088266 | Ta_SA5_6h - Ta_CTRL_0h | 2.66E-03 | 0.707 | -0.738 |
| LOC123132638 | Uncharacterized protein LOC123132638 | Ta_SA5_6h - Ta_CTRL_0h | 2.66E-03 | 0.321 | -0.219 |
| LOC123078822 | Uncharacterized protein LOC123078822 | Ta_SAMv2_6h - Ta_SA5_6h | 2.67E-03 | 0.606 | -0.338 |
| LOC123138436 | Uncharacterized LOC123138436 | Ta_SAMv2_6h - Ta_SA5_6h | 2.67E-03 | 0.867 | -0.171 |
| LOC123091552 | Uncharacterized LOC123091552 | Ta_SA1_6h - Ta_ECC_6h | 2.68E-03 | -1.000 | -0.074 |
| LOC123103571 | ASC1-like protein 1 | Ta_SAMv2_6h - Ta_SA1_6h | 2.76E-03 | -0.587 | 0.401 |
| LOC123156386 | Uncharacterized LOC123156386 | Ta_SAMv2_6h - Ta_SA1_6h | 2.76E-03 | -0.577 | -0.201 |
| LOC542897 | E3 ubiquitin-protein ligase RHF2A | Ta_SAMv2_6h - Ta_SA1_6h | 2.76E-03 | -0.207 | -0.106 |
| LOC123093693 | Uncharacterized protein LOC123093693 | Ta_SAMv2_6h - Ta_SA1_6h | 2.76E-03 | -0.902 | 0.088 |
| LOC123088517 | Uncharacterized protein LOC123088517 | Ta_SAMv2_6h - Ta_SA1_6h | 2.76E-03 | 0.375 | 0.004 |
| LOC123079583 | Probable serine/threonine-protein kinase PBL1 | Ta_SAMv2_6h - Ta_ECC_6h | 2.90E-03 | 0.966 | 0.605 |
| LOC123091823 | Uncharacterized LOC123091823 | Ta_SAMv2_6h - Ta_ECC_6h | 3.25E-03 | -0.413 | -0.473 |
| LOC123116107 | Uncharacterized protein LOC123116107 | Ta_SAMv2_6h - Ta_SA1_6h | 3.56E-03 | 1.000 | 0.623 |
| LOC123138430 | Transcription factor TFIIIB component B''-like | Ta_SA1_6h - Ta_SA5_6h | 3.57E-03 | -0.952 | 0.222 |
| LOC123137370 | Uncharacterized LOC123137370 | Ta_SA1_6h - Ta_SA5_6h | 3.57E-03 | 0.969 | 0.015 |
| LOC123051132 | Uncharacterized protein LOC123051132 | Ta_SA1_6h - Ta_SA5_6h | 3.83E-03 | -1.000 | 1.027 |
| LOC123063450 | Crossover junction endonuclease MUS81-like | Ta_SA1_6h - Ta_SA5_6h | 3.83E-03 | -1.000 | -0.155 |
| LOC123046508 | Cation transporter HKT4-like | Ta_SAMv2_6h - Ta_CTRL_0h | 4.28E-03 | 0.759 | -1.862 |
| LOC123045258 | Phospholipid-transporting ATPase 2-like | Ta_SAMv2_6h - Ta_CTRL_0h | 4.28E-03 | 1.000 | 0.792 |
| LOC123146363 | Protein argonaute 1C-like | Ta_SAMv2_6h - Ta_CTRL_0h | 4.28E-03 | -0.276 | -0.176 |
| LOC123123592 | Uncharacterized LOC123123592 | Ta_SA5_6h - Ta_ECC_6h | 4.55E-03 | -0.800 | 0.003 |
| LOC123185486 | Monooxygenase 2-like | Ta_SAMv2_6h - Ta_CTRL_0h | 4.62E-03 | 0.874 | 0.067 |
| LOC123070240 | Uncharacterized LOC123070240 | Ta_SA5_6h - Ta_ECC_6h | 4.75E-03 | -0.613 | -0.456 |
| LOC123149503 | Uncharacterized LOC123149503 | Ta_SA5_6h - Ta_ECC_6h | 4.75E-03 | 0.241 | -0.122 |
| LOC123122996 | Pentatricopeptide repeat-containing protein At2g22070-like | Ta_SA5_6h - Ta_ECC_6h | 4.75E-03 | 1.000 | -0.068 |

| | | | | | |
|---|---|---|---|---|---|
| LOC123138430 | Transcription factor TFIIIB component B''-like | Ta_SA1_6h - Ta_ECC_6h | 5.23E-03 | -0.796 | 0.327 |

**Table S12**. Plant Reactome pathways significantly enriched with *Triticum aestivum* differentially expressed genes from all contrasts to the 0 h control (Ta_ECC_6h - Ta_CTRL_0h, Ta_SA1_6h - Ta_CTRL_0h, Ta_SAMv2_6h - Ta_CTRL_0h, Ta_SA5_6h - Ta_CTRL_0h). Protein blast IDs and Gene Ontology were linked to Plant Reactome pathways. Presented *p*-values were obtained by performing pathway enrichment analyses using Fisher's test followed by adjustment for FDR. Highlighted *p*-values > 0.05.

| | **FDR-adjusted *p*-value** | | | |
|---|---|---|---|---|
| **Plant Reactome pathway** | Ta_ECC_6h - Ta_CTRL_0h | Ta_**SA1**_6h - Ta_CTRL_0h | Ta_**SAMv2**_6h - Ta_CTRL_0h | Ta_**SA5**_6h - Ta_CTRL_0h |
| Oleoresin sesquiterpene volatiles biosynthesis | 1.0000 | **0.0000** | **0.0037** | **0.0000** |
| Monoterpene biosynthesis | 1.0000 | **0.0021** | 1.0000 | 0.1669 |
| Thiamine biosynthesis | **0.0282** | 1.0000 | **0.0090** | **0.0034** |
| Jasmonic acid biosynthesis | 1.0000 | 0.3276 | **0.0037** | 0.0528 |
| Photorespiration | 1.0000 | 1.0000 | **0.0037** | 1.0000 |
| Gibberellin biosynthesis I | **0.0206** | 1.0000 | **0.0077** | **0.0226** |
| Methylerythritol phosphate pathway | 0.7613 | 0.0620 | **0.0325** | **0.0127** |
| Abscisic acid mediated signalling | 0.5401 | **0.0491** | 0.2367 | **0.0169** |
| Trans, trans-farnesyl diphosphate biosynthesis | 1.0000 | 0.0898 | 0.0574 | **0.0226** |
| Flavin biosynthesis | 1.0000 | **0.0281** | 0.5849 | 0.4477 |
